# RHEB/mTOR-hyperactivity causing cortical malformations drives seizures through increased axonal connectivity

**DOI:** 10.1101/2020.07.08.189399

**Authors:** Martina Proietti Onori, Linda M.C. Koene, Carmen B. Schafer, Mark Nellist, Marcel de Brito van Velze, Zhenyu Gao, Ype Elgersma, Geeske M. van Woerden

**Author notes:** These authors contributed equally (Linda M.C. Koene, Carmen B. Schafer). Correspondence (Y.E.), (G.M.v.W.).

## Abstract

Dominant-active mutations in *Ras Homolog Enriched in Brain 1* (*RHEB*), such as the recently identified RHEBp.P37L mutation, can cause malformations of cortical development (MCD) with associated epilepsy and intellectual disability through a yet unknown mechanism. We found that focal expression of RHEBp.P37L in mouse somatosensory cortex results in an MCD-like phenotype, with increased mammalian target of rapamycin (mTOR) signaling, ectopic localization of neurons and generalized seizures. In addition, the RHEBp.P37L expressing neurons showed increased axonal length and branching. By temporally controlling RHEBp.P37L expression, we found that the cortical malformation by itself was neither necessary nor sufficient to generate seizures. Rather, seizures were contingent on persistent mTOR activation and enhanced axonal connectivity of RHEBp.P37L expressing neurons, causing hyperexcitability of distally connected neurons. These results provide new evidence of the extent of anatomical and physiological abnormalities caused by mTOR hyperactivity, beyond local malformations, that can lead to generalized epilepsy.

## INTRODUCTION

Malformations of cortical development (MCD) are a heterogenous group of micro- and macroscopic cortical abnormalities, such as focal cortical dysplasia (FCD), megalencephaly, lissencephaly and periventricular nodular heterotopia (Barkovich et al., 2012). MCD arise from disturbances in cortical development during early embryogenesis and are often linked to epilepsy and intellectual disability (ID) (Juric-Sekhar and Hevner, 2019; Leventer et al., 2008; Represa, 2019). It is estimated that up to 40% of intractable or difficult to control childhood seizures are due to MCD, and *vice versa*, at least 75% of the patients with MCD will develop seizures (Leventer et al., 1999).

The mammalian (or mechanistic) target of rapamycin (mTOR) is a kinase that mediates many cellular processes, including neuronal progenitors proliferation and cell growth (Laplante and Sabatini, 2012; Saxton and Sabatini, 2017). mTOR forms 2 distinct protein complexes, characterized by different binding partners, mTORC1 and mTORC2 (Bhaskar and Hay, 2007). mTORC1 is regulated by the tuberous sclerosis complex (TSC) and the Ras Homolog Enriched in Brain 1 (RHEB) (Parmar and Tamanoi, 2010). RHEB, a member of the RAS family of small GTPases, is the direct activator of mTORC1 (Bai et al., 2007). The conversion of active GTP-bound RHEB to the inactive GDP-bound form is mediated by the TSC complex, which acts as a RHEB GTPase activating protein (GAP) (Li et al., 2004). In response to nutrients and growth factors the TSC complex is inhibited, allowing activation of mTORC1 by RHEB-GTP (Manning and Cantley, 2003; Sabatini, 2017). Studies in *Rheb* knock-out mice showed that RHEB activity is the rate limiting step for mTOR activation in the brain, and that neuronal functioning in particular is sensitive to increased RHEB-mTOR signaling (Goorden et al., 2015).

Hyperactivation of the mTOR pathway by mutations in genes encoding components of the mTOR pathway (*e.g. AKT3*, *PIK3CA*, *DEPDC5*, *PTEN*, *TSC1, TSC2, RHEB* and *MTOR* itself) has been associated with different types of MCD, such as megalencephaly and FCD, as well as with epilepsy (Crino, 2011; Juric-Sekhar and Hevner, 2019; Moffat et al., 2015). The underlying genetic variability explains the heterogeneity of MCD and illustrates the challenges involved in understanding the mechanisms underlying MCD-associated epilepsy.

The discovery of genetic mutations that cause FCD or other types of MCD, allowed the generation of animal models to study the development of MCD and associated epilepsy (Chevassus-au-Louis et al., 1999; Wong and Roper, 2016). In particular, *in utero* electroporation (IUE), that allows for the spatial and temporal control of transgene expression during embryonic development, has been used to generate mouse models with focal malformations and epilepsy (Hanai et al., 2017; Hsieh et al., 2016; Park et al., 2018; Ribierre et al., 2018).

One recent FCD mouse model was generated by using IUE to overexpress the constitutively active RHEBp.S16H mutant (Yan et al., 2006). This results in mTOR hyperactivity, FCD and spontaneous seizures (Hsieh et al., 2016). Recently we identified two *de novo* mutations in *RHEB* (c.110C>T (p.P37L) and c.202T>C (p.S68P)) in patients with ID, epilepsy and megalencephaly (Reijnders et al., 2017), providing for the first time a clinically relevant link between RHEB and MCD. IUE of a construct encoding the RHEBp.P37L mutant caused severe focal cortical lesions, resembling periventricular nodular heterotopia, and diffuse neuronal misplacement in the cortex. Furthermore, the mice reliably developed spontaneous seizures starting at three weeks of age (Reijnders et al., 2017). The anatomical and phenotypical features of this novel mouse model, fully recapitulating the most prominent characteristics of MCD (focal lesions and epilepsy), make this a powerful tool and clinically relevant novel model to study the mechanisms underlying mTOR and MCD-related epilepsy.

Here we demonstrate that *RHEB* mutations that cause MCD in patients activate mTORC1 signaling and that the cortical malformation is not required for the development of mTOR related seizures, in line with previous studies (Abs et al., 2013; Hsieh et al., 2016). Additionally, we provide evidence that the presence of heterotopia by itself is insufficient to cause epilepsy. Instead, we found that persistent activation of the mTOR pathway results in anatomical and functional changes in axonal connectivity, which cause increased excitability of distally connected neurons and the development of generalized seizures.

## RESULTS

### The RHEBp.P37L protein is resistant to TSC complex inhibition and causes aberrant cortical development *in vivo*

The RHEBp.P37L mutation was identified in patients with ID, megalencephaly and epilepsy, and it was proposed to act as a gain of function mutation (Reijnders et al., 2017). This could potentially be due to resistance to the GAP-function of the TSC complex. To assess whether the TSC complex can convert RHEBp.P37L from its active GTP- to its inactive GDP-bound state, we compared the effects of transient *in vitro* overexpression of the RHEBp.P37L mutant with wild-type RHEB (RHEB WT) and the RHEBp.S16H mutant, a well-known gain of function mutant of RHEB recently used to generate an FCD mouse model (Hsieh et al., 2016; Yan et al., 2006). In the absence of TSC, overexpression of RHEB WT as well as both RHEB mutants caused increased mTORC1 activity, as measured by T389-phosphorylation of co-expressed S6K, a direct substrate of the mTORC1 kinase (Figure 1A, see **Supplementary table1** for statistics overview). In the presence of the TSC complex, mTORC1 activity was reduced in the RHEB WT and RHEBp.S16H expressing cells, but not in the RHEBp.P37L expressing cells (Figure 1A). Here, presence or absence of TSC resulted in similar levels of S6K phosphorylation, confirming that the patient-derived RHEBp.P37L acts as a gain of function mutation which is resistant to inhibitory action of the TSC complex (Figure 1A).

**Figure 1.**
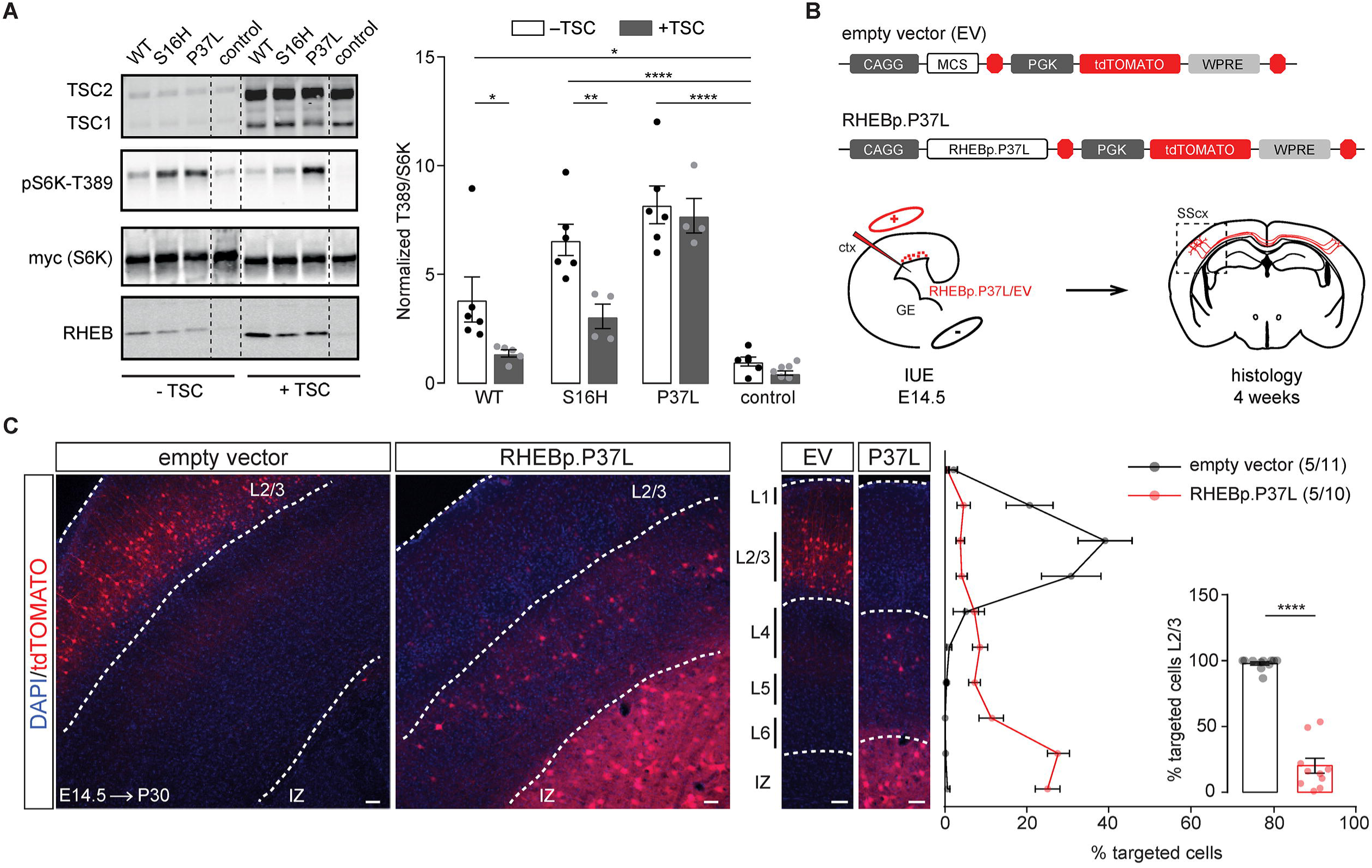
The RHEBp.P37L protein is resistant to TSC complex inhibition and causes aberrant cortical development *in vivo*. **(A)** Wild-type RHEB (WT), p.S16H or p.P37L constructs were transiently co-expressed with an S6K reporter construct and the TSC complex in HEK 293T cells to assess the effect on mTORC1 activity. Quantification of the ratio of T389-phosphorylated S6K to total S6K was calculated relative to control condition, in absence or presence of the TSC complex (control indicates empty vector pcDNA3); dashed lines indicate where an irrelevant lane in the original scan was excluded from the picture; bar graph represents mean ± SEM and single data points represent the number of independent biological samples per condition; for statistics see **Supplementary table 1**. **(B)** Schematic representation of the main constructs used throughout the experiments and overview of the experimental design. MCS indicate the multiple cloning site with specific restriction sites (AscI and PacI in this case) to insert the gene of interest. Each construct was delivered by *in utero* electroporation (IUE) at E14.5 to target the progenitor cells of layer 2/3 pyramidal neurons in the somatosensory cortex (SScx); ctx=cortex; GE=ganglion eminence. **(C)** Representative confocal images of the targeted SScx counterstained with DAPI showing the transfected cells (tdTomato+) in red (see also **Figure 1-figure supplement 1**) and quantification of tdTomato+ cells across the different layers of the SScx with percentage of cells reaching layer 2/3 (L2/3) in the inset (bins 1-5 from the top). Dotted lines indicate the border of the intermediate zone (IZ, bottom) and delineate L2/3. Numbers in the legend indicate number of targeted mice (N=5) and total number of pictures analyzed (n=11, n=10); results are represented as mean ± SEM and single data points in the bar graph indicate the number of pictures analyzed; inset analysis: Mann-Whitney U = 0, *p*<0.0001, two-tailed Mann-Whitney test. * *p*<0.05, ** *p*<0.01, **** *p*<0.0001; scale bars: 50 μm.

Using IUE, we have previously shown that overexpression of the RHEBp.P37L mutant, but not RHEB WT, results in the formation of a heterotopic nodule as well as spontaneous epilepsy in 100% of the targeted mice (Reijnders et al., 2017), providing us with a valuable model to study the mechanisms behind mTORC1-dependent and MCD-related epilepsy. To confirm previous results and further characterize the model, we used IUE to introduce the RHEBp.P37L vector or an empty vector control at E14.5 in progenitor cells that give rise to layer 2/3 (L2/3) pyramidal neurons of the somatosensory cortex (SScx) (Figure 1B). As shown previously, overexpression of RHEBp.P37L resulted in a clear migration deficit, with only 20% of the targeted cells reaching the outer layers of the cortex (L2/3) compared to 97% in the empty vector condition (Figure 1C, inset). The non-migrated transfected neurons remained in the white matter to form a heterotopic nodule lining the ventricle in the adult brain (Figure 1C and **Figure 1-figure supplement 1**).

While the general cortical layer architecture remained intact (**Figure 1-figure supplement 1**, ectopic RHEBp.P37L overexpressing cells showed cytological abnormalities, with dysmorphic appearance and enlarged soma size (Figure 2A and **Figure 1-figure supplement 1**). Sholl analysis of biocytin filled cells in the heterotopic nodule of RHEBp.P37L expressing neurons revealed that the cells in the nodule presented a more complex arborization compared to empty vector control cells in L2/3 (Figure 2B). Transfected ectopic neurons preserved the molecular identity of mature L2/3 pyramidal cells, being positive for the neuronal marker NeuN and the outer layer molecular marker CUX1 and negative for the deeper layer marker CTIP2 (Figure 2C and **Figure 2-figure supplement 1A**). Additionally, most neurons in the nodule were SATB2 positive, showing that, despite being mislocalized, they maintained the callosal projection identity (Figure 2C and **Figure 2-figure supplement 1A**). Finally, mice overexpressing RHEBp.P37L showed an overall increase in ribosomal protein S6 phosphorylation, a commonly used readout for mTORC1 activity, in the transfected hemisphere compared to the empty vector condition (Figure 2D and **Figure 2-figure supplement 1B**).

**Figure 2.**
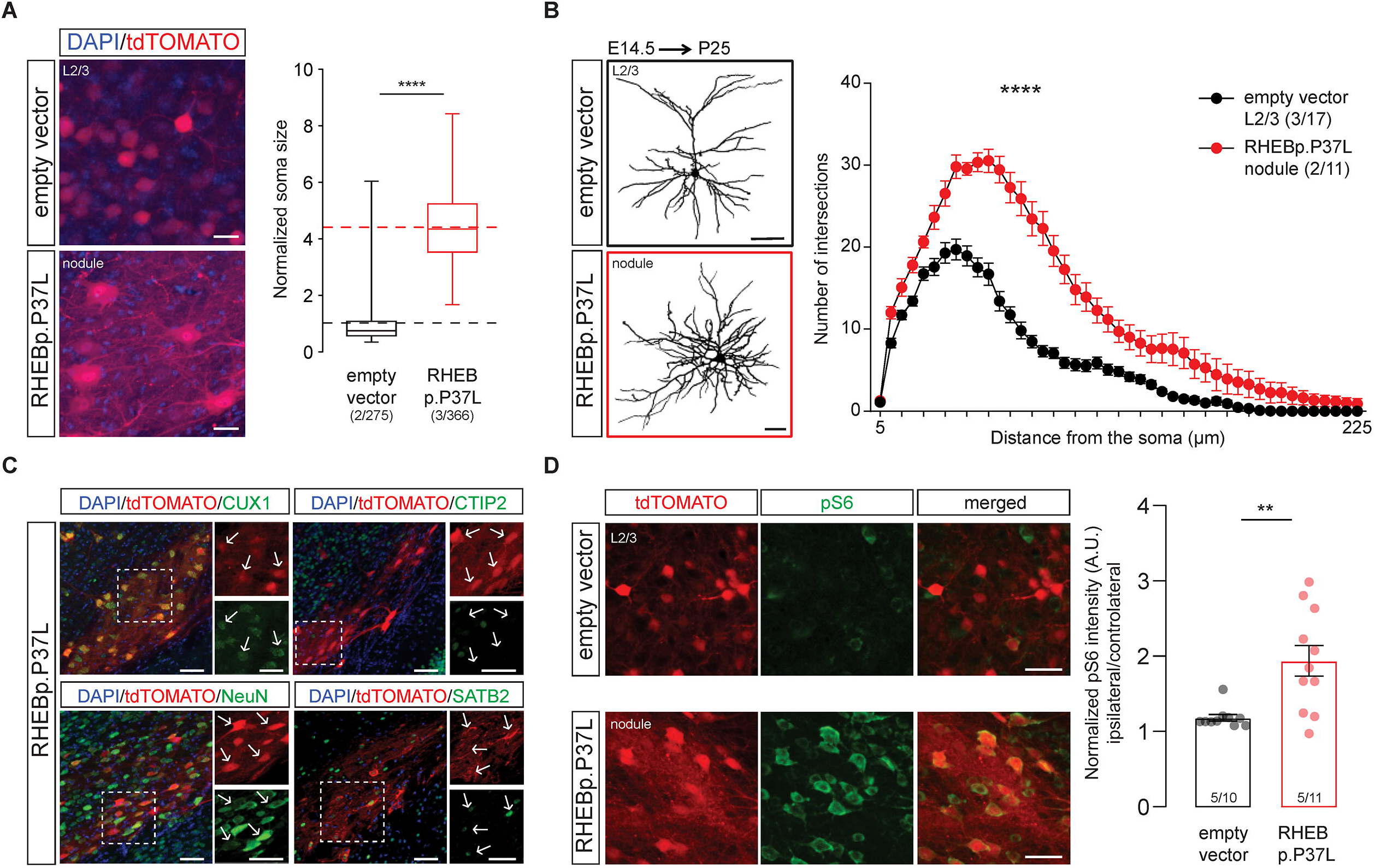
Ectopic RHEBp.P37L cells display aberrant morphology and show mTOR hyperactivity while still preserving the molecular identity of L2/3 neurons. **(A)** Soma size quantification of L2/3 empty vector expressing cells and RHEBp.P37L expressing cells in the nodule; box plots represent minimum and maximum values with median, dashed lines represent the mean values for empty vector (black) and for RHEBp.P37L (red); numbers indicate number of targeted mice (N=2, N=3) and number of cells analyzed (n=275, n=366); Mann-Whitney U = 1940, *p*<0.0001, two-tailed Mann-Whitney test. **(B)** Reconstruction and Sholl analysis of dendritic morphology of biocytin filled cells in L2/3 of the SScx (for empty vector control) and RHEBp.P37L cells in the nodule; numbers in the legend indicate number of targeted mice (N=3, N=2) and number of cells analyzed (n=17, n=11); data are presented as mean ± SEM; interaction group condition/distance from the soma: F(44, 1144) = 15.69, mixed-effects analysis; *p*<0.0001. **(C)** Representative images of the nodule stained with CUX1 (L2/3 marker), CTIP2 (L5 marker), SATB2 (cortical projection neurons marker) or NeuN (mature neurons marker); arrows in the zoomed pictures point at examples of targeted cells; for an overview see **Figure 2-figure supplement 1**. **(D)** Representative images of the targeted L2/3 (SScx) of empty vector control and nodule showing increased pS6-240 levels for the ipsilateral targeted cortex in RHEBp.P37L targeted mice; for an overview see **Figure 2-figure supplement 1**; bar graph represents mean ± SEM and single data points indicate the values of each normalized ipsilateral/contralateral pS6 intensity; numbers in the bars indicate number of targeted mice (N=5) and number of pictures analyzed (n=10, n=11); Mann-Whitney U = 13, *p* = 0.002, two-tailed Mann-Whitney test). Histological analysis for **(A) (C)** and **(D)** was performed on 5 weeks old mice. ** *p*<0.01, **** *p*<0.0001; scale bars: 20 μm (A), 50 μm (B-D).

### Overexpression of RHEBp.P37L *in vivo* causes mTORC1-dependent spontaneous generalized tonic-clonic seizures and abnormal neuronal network activity

To assess the reliability of spontaneous seizures development, the RHEBp.P37L mice were continuously monitored from weaning (P21) using wireless electroencephalography (EEG) (Figure 3A). Spontaneous seizures started to appear at 3 weeks of age, with an average onset of 33 days, confirming previous data (Reijnders et al., 2017) (**Figure 3-figure supplement 1A**). Seizures were highly stereotypical, characterized by the loss of upright posture followed by a tonic-clonic phase with convulsions and twitching behavior. EEG recordings showed that, while control mice did not show any epileptic activity (N = 6), all RHEBp.P37L mice (N = 12) showed clear epileptic events (Figure 3B and **Figure 3-figure supplement 1B**). Seizures were characterized by an increase in frequency and amplitude of brain activity (Figure 3C, box 3 ictal activity) compared to baseline interictal activity (Figure 3C, box 2) and baseline activity of control mice (Figure 3C, box 1). The calculated average duration of an epileptic event was 40 seconds (mean ± SEM: 42.6 ± 1.33), followed by a post-ictal depression phase of variable length (Figure 3C, box 4 post-ictal activity). The frequency of seizures per day was variable between mice as well as per mouse over time (**Figure 3-figure supplement 1C**). Additionally, no correlation was found between the total number of seizures over three consecutive days of recording and the average number of targeted cells per mouse (**Figure 3-figure supplement 1D**).

**Figure 3.**
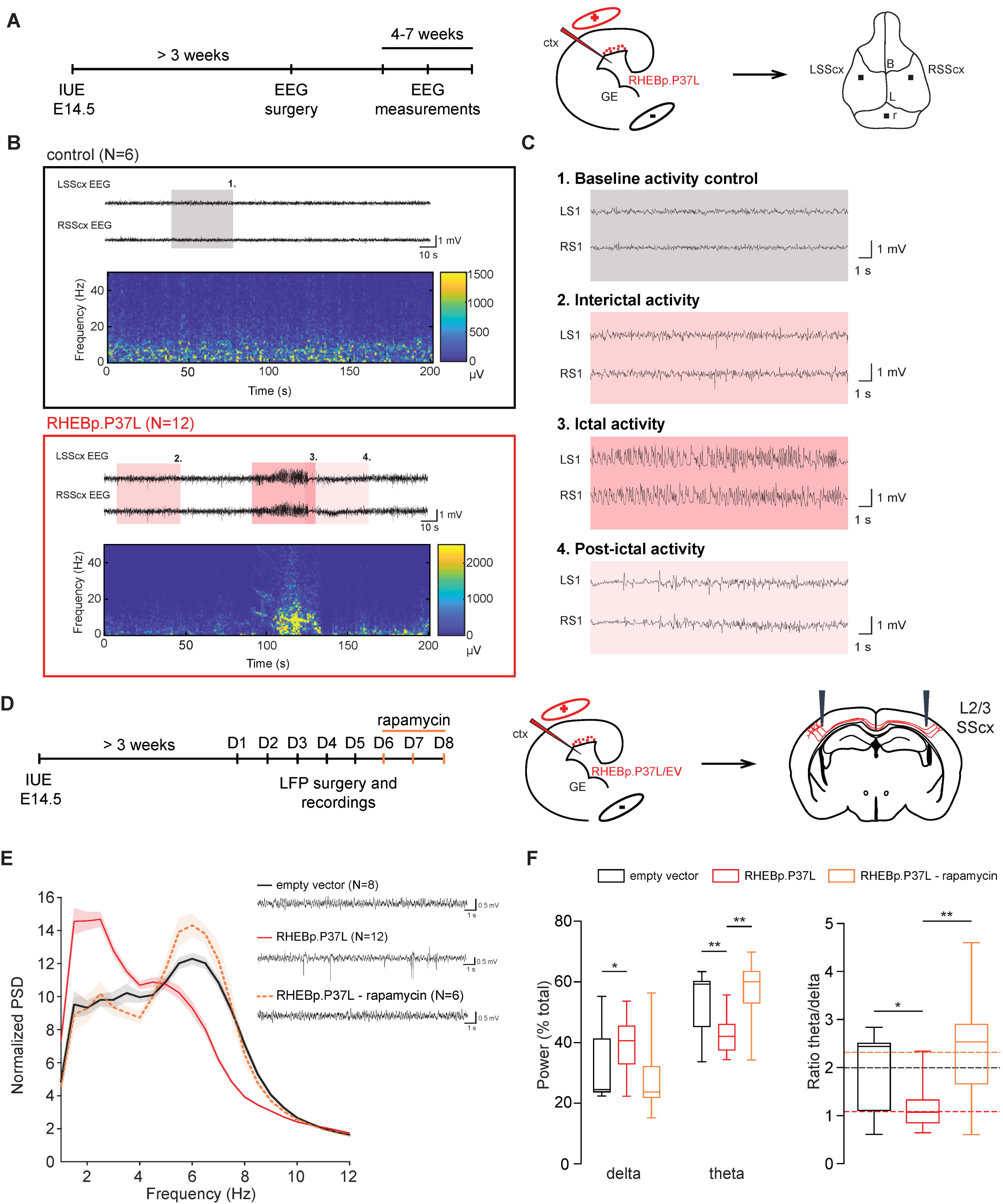
Overexpression of RHEBp.P37L *in vivo* causes mTORC1-dependent spontaneous generalized tonic-clonic seizures and abnormal neuronal network activity. **(A)** Timeline and experimental design indicating the cortical area targeted with the IUE and position of the electrodes placed during the EEG surgery (LSScx = left SScx; RSScx = right SScx; B = bregma; L= lambda; r = reference electrode). **(B)** Example EEG traces and spectrogram of 5 weeks old control mouse (N=6, non-targeted mice from the same litters as the RHEBp.P37L mice) and RHEBp.P37L mouse (N=12); see also **Figure 3-figure supplement 1**; colored boxes are zoomed in panel **(C)**. **(C)** Highlighted EEG traces showing: box 1. the baseline activity of a control mouse; box 2. the interictal activity, box 3. the ictal (seizure) activity and box 4. the post-ictal phase of a RHEBp.P37L targeted mouse. **(D)** Timeline and experimental design indicating the cortical area targeted with the IUE, the position of the electrodes for the local field potential (LFP) recordings and the IP rapamycin injections. **(E)** Example LFP traces for each group condition and normalized power spectrum density (PSD) averaged bilaterally over the overall consecutive days of recording (for the PSD until 50 Hz see **Figure 3-figure supplement 1**); N indicates number of mice analyzed for each group; data are represented as mean (thick line) ± SEM (shading area). **(F)** Calculation of the *delta* (2-4 Hz) and *theta* (4-8 Hz) frequency bands over the total power of the PSD presented in **(E)**, and relative ratio *theta*/*delta* (see also **Figure 3-figure supplement 1**); box plots represent minimum and maximum values with median, dashed lines represent the mean values for each group; for statistics see **Supplementary table 2**; * *p*<0.05, ** *p*<0.01, *** *p*<0.001.

To assess if brain-wide suppression of mTORC1 activity could reduce seizures, we treated a group of mice showing seizures (N=6; 5-6 weeks old) systemically for 7 days with the allosteric mTORC1 inhibitor rapamycin (10 mg/kg). Rapamycin treatment reduced and temporarily abolished the occurrence of seizures within one week from the last day of rapamycin administration. However, seizures reoccurred starting 3 weeks after the last injection of rapamycin in 4 out of 6 mice, indicating that sustained inhibition of mTORC1 is required to fully suppress the seizures (data not shown).

Electrographic frequency dynamics of the interictal phases, and especially *theta* oscillations, have been proven to be good predictors for epilepsy outcome, compared to epileptiform spikes or high-frequency oscillations (HFOs), in several rodent models of epilepsy (Chauvière et al., 2009; Milikovsky et al., 2017). Therefore, using local field potential (LFP) recordings, we assessed the frequency dynamics of cortical brain activity in the interictal periods of RHEBp.P37L expressing mice, starting from 4 weeks of age (Figure 3D). The normalized averaged power spectrum of the RHEBp.P37L group did not reveal a significant difference between the targeted and non-targeted cortex (targeting: F(1, 22)=1.43, p=0.25, non-significant, Two-way repeated measure ANOVA; data not shown), therefore measurements from both sides were pooled. Whereas the total power across 5 days of recording did not differ between the RHEBp.P37L (N=12) and the control group (N=8) (Mann-Whitney U = 157, p = 0.35, non-significant, two-tailed Mann-Whitney test, data not shown), a significant difference in the *delta* (2-4 Hz), *theta* (4-8 Hz) and *gamma* (30-50 Hz) frequency bands of the normalized power spectrum was seen in the RHEBp.P37L group compared to the control group (Figure 3E, 3F and **Figure 3-figure supplement 1E and 1F**; statistics in **Supplementary table2**). The difference in the *theta* and *gamma* frequency bands, but not in the *delta*, could be reverted to the control condition by injecting the RHEB mice with 10 mg/kg rapamycin intraperitoneally for 3 consecutive days (Figure 3E, 3F and **Figure 3-figure supplement 1E and 1F**; statistics in **Supplementary table2**). Together with the finding that rapamycin abolished seizures, this result indicates that *theta* oscillations, which negatively correlate with *gamma* frequencies (Milikovsky et al., 2017), are a good predictor for epileptogenesis in the RHEBp.P37L mouse model.

### The heterotopic nodule is neither necessary nor sufficient to induce spontaneous seizures

Cortical malformations occur during early embryonic development and are generally associated with the development of epileptic activity (Represa, 2019). Therefore, a transient treatment with mTOR inhibitors during brain development might prevent the formation of a cortical malformation and could consequently reduce the chances of developing epilepsy. To assess if early transient down-regulation of the mTORC1 pathway upon overexpression of RHEBp.P37L could prevent the development of heterotopic nodules, we injected pregnant female mice with 1 mg/kg of rapamycin for 2 consecutive days starting 1 day after IUE of the RHEBp.P37L vector (Figure 4A). Prenatal down-regulation of the mTORC1 pathway significantly improved the migration of the targeted neurons, with 75% of the targeted cells successfully migrating out (Figure 4B). In addition, prenatal rapamycin treatment successfully prevented the formation of a heterotopic nodule in 9 out of 11 mice. However, 7 out of the 11 targeted mice (58%) still showed spontaneous seizures, including 5 mice that did not develop a discernable heterotopic nodule (Figure 4C). Average onset of seizures was comparable to the non-treated RHEBp.P37L mice (mean ± SEM: 32.6 days ± 2.3; Chi square (1) = 0.16, *p* = 0.69, Log-rank test, data not shown). Hence, the presence of a heterotopic nodule is not required for RHEBp.P37L mediated seizures, and reducing the formation of these nodules does not always prevent epileptogenesis.

**Figure 4.**
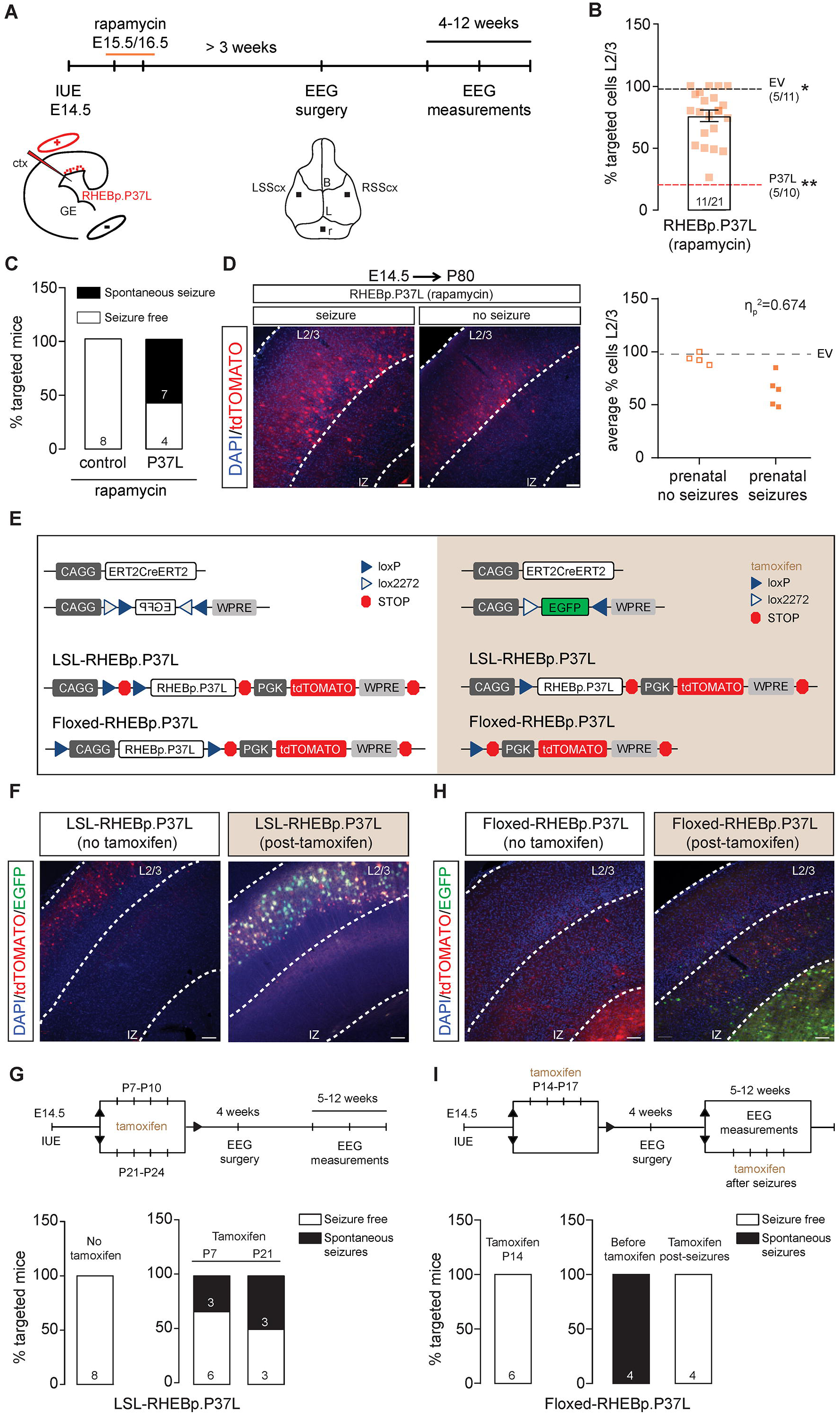
The heterotopic nodule is neither necessary nor sufficient to induce spontaneous seizures. **(A)** Schematic representation of the timeline of the IUE, SC rapamycin injections, EEG surgery and measurements. **(B)** Quantification of the percentage of tdTomato+ cells that managed to migrate out to L2/3 in mice prenatally exposed to rapamycin; data are presented as mean ± SEM, single data points represent the values of each picture analyzed and dashed lines indicate the mean value of cells reaching L2/3 in empty vector control mice (black line) and in RHEBp.P37L mice (red line); numbers in the graph indicate number of mice (N=11, N=5) and number of pictures analyzed (n=21, n=11, n=10); (% targeted cells in L2/3: H(2) = 25.97, *p* < 0.0001, Kruskal-Wallis test; EV *vs* RHEBp.P37L-prenatal rapamycin, *p* = 0.05; RHEBp.P37L *vs* RHEBp.P37L-prenatal rapamycin, *p* = 0.002, RHEBp.P37L *vs* EV, *p* < 0.0001, Dunn’s multiple comparisons test; * *p*<0.05, ** *p*<0.01. **(C)** Percentage of targeted mice showing spontaneous seizures; control mice are non-targeted mice from the same litters as the RHEBp.P37L mice prenatally exposed to rapamycin; numbers in the bar plots indicate the number of mice per group. **(D)** Representative images of RHEBp.P37L mice prenatally exposed to rapamycin that showed or did not show seizures and degree of association between the migration phenotype (mean value of % of targeted cells in L2/3, dependent scale variable, for each mouse shown in figure B) and the absence or presence of seizures (independent nominal variable) in RHEBp.P37L mice (N=4 and N=5, respectively, with the exclusion of the mice that showed heterotopia); the dashed line represents the mean value of the empty vector control group already shown in B, as comparison; η_p_=0.821, η_p_ =0.674, Eta measure of association, with values of η_p_ close to one indicating strong association. **(E)** Schematic representation of the DNA plasmids used in the experiment. The Lox-Stop-Lox (LSL) or the floxed construct was expressed in combination with the CAGG-ERT2CreERT2 and a CAGG-DIO-EGFP constructs. The EGFP in the CAGG-DIO-EGFP construct is expressed only upon tamoxifen injection, providing a measure of efficient cre-dependent recombination. **(F)** and **(H)** Representative images showing the effect of tamoxifen administration in adult mice injected *in utero* with either the LSL construct **(F)** or the floxed construct **(G)**; mice were injected 4 times with tamoxifen starting from P7 in **(F)** and starting from P14 in **(G)** and sacrificed at P50. **(G)** and **(I)** Timeline of the experimental design and bar graphs indicating the percentage of targeted mice for each group measured with EEG until 12 weeks of age showing spontaneous seizures; numbers in the bar plots indicate the number of mice. Scale bars: 100 μm.

When comparing the cortical migration patterns in mice with and without seizures, a clear correlation was observed between the migration pattern of RHEBp.P37L expressing cells and the presence or absence of seizures: RHEBp.P37L-prenatal treated mice with seizures showed a more severe migration deficit of RHEBp.P37L expressing cells compared to prenatal treated RHEBp.P37L expressing mice that were seizure free (Figure 4D). In fact, the percentage of cells that reached L2/3 of the SScx of RHEBp.P37L-prenatal treated mice with seizures (63%), was significantly lower than RHEBp.P37L-prenatal treated mice without seizures (93%) or control mice (98%) (% targeted cells in L2/3: H(2) = 22.08, *p* < 0.0001, Kruskal-Wallis test; empty vector *vs* RHEBp.P37L-no seizures, *p* > 0.99; empty vector *vs* RHEBp.P37L-seizures, *p* < 0.0001; RHEBp.P37L-no seizures *vs* RHEBp.P37L-seizures, *p* = 0.002; Dunn’s multiple comparisons test, data not shown). These results indicate that ectopic cells do facilitate the process of epileptogenesis.

Hyperactivation of mTORC1 is sufficient to cause seizures, independent of the presence of cortical malformations (Abs et al., 2013), even when the mTORC1 activity is increased in a limited set of neurons (Hsieh et al., 2016). Moreover, the cortical malformation by itself, in the absence of continued mTORC1 signaling, does not cause epilepsy, as was also shown by brain-wide inhibition of mTORC1signaling (Hsieh et al., 2016). To further dissect the relationship between increased mTOR activity, cortical malformations and epilepsy, we made use of a genetic approach, that allowed us not only to regulate mTORC1 activity in a temporal fashion, but also to restrict its activity to a limited number of cells. To that end we used IUE to introduce a Lox-Stop-Lox(LSL)-RHEBp.P37Lvector or floxed-RHEBp.P37L vector together with a vector expressing the ERT2CreERT2 fusion protein (Figure 4E). This allowed us to switch the RHEBp.P37L gene respectively on or off during different stages of cortical development by means of systemic tamoxifen administration. The use of an additional vector that expresses EGFP in a Cre-dependent manner (CAG-DIO-EGFP), allowed us to assess the efficiency of Cre activation upon tamoxifen administration. IUE of the LSL-RHEBp.P37L construct in the absence of tamoxifen administration, did not result in a migration deficit, or seizures. This indicates that the LSL cassette successfully prevented RHEBp.P37L expression (Figure 4F). However, once expression of RHEBp.P37L was induced by administration of tamoxifen either at P7 or P21, a subset of the mutant mice (38% of the P7 group and 50% of the P21 group) developed spontaneous seizures (Figure 4G), albeit with a delayed onset compared to mice that express RHEBp.P37L throughout development (Chi square (2) = 10.18, *p* =0.006; Log-rank test, data not shown). This indicates that RHEBp.P37L expression in a limited number of cells, can drive seizures in the absence of an observable cortical malformation (migration defects or heterotopic nodules).

To investigate if presence of the heterotopic nodule is epileptogenic after normalizing mTORC1 activation only in the targeted cells (instead of brain-wide as others also did in previous studies (Hsieh et al., 2016)), we used IUE to insert the floxed-RHEBp.P37L vector (Figure 4H). Upon expression of RHEBp.P37L during early pre- and postnatal development, we normalized mTORC1 activity in these cells by inducing RHEBp.P37L deletion at P14. (Figure 4I). Although a clear heterotopic nodule was formed in these mice, none of the mice developed seizures (Figure 4I). Furthermore, inducing deletion of RHEBp.P37L after epileptogenesis, completely abolished the seizures within 10 days from gene deletion (N=4, last EEG measurements performed between day 85 and 90) (Figure 4I). Taken together, these results indicate that cortical malformations are neither necessary nor sufficient for the development of spontaneous seizures in our mouse model.

### RHEBp.P37L expression induces aberrant axonal development both *in vitro* and *in vivo* and functional increased contralateral L2/3 and L5 connections

The mTOR pathway plays an important role in axonal outgrowth, with functional effects on neuronal network formation (Choi et al., 2008; Gong et al., 2015; Nie et al., 2010). Because increasing mTOR signaling in a limited number of neurons in the brain is enough to cause seizures, independently from cell misplacement, we hypothesized that this could be due to aberrant neuronal connectivity caused by RHEBp.P37L overexpression. Therefore, we investigated the effect of RHEBp.P37L on axonal length and branching. Overexpression of RHEBp.P37L in primary hippocampal neurons *in vitro* caused a significant increase in axonal length and axonal branching, compared to the empty vector control (Figure 5A).

**Figure 5.**
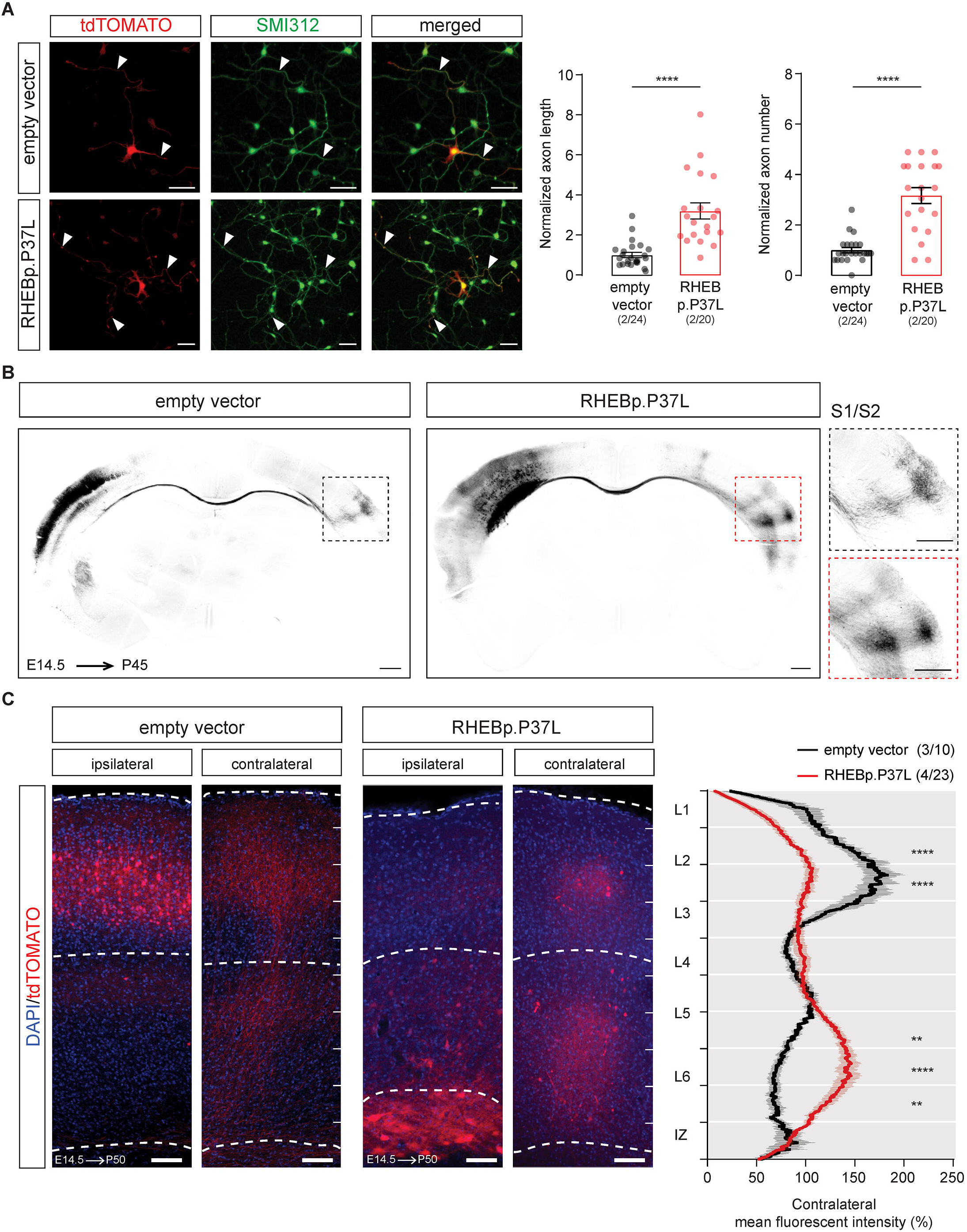
RHEBp.P37L overexpression induces an increase in axon length and branching both *in vitro* and *in vivo*. **(A)** Representative images of primary hippocampal cultures transfected at day *in vitro* 1 (DIV1) with either empty vector control or RHEBp.P37L constructs (tdTomato, in red) stained at DIV4 with a pan axonal marker SMI312 (in green); arrowheads indicate the axons; bar graphs represent mean ± SEM and single data points indicate the number of cells analyzed; numbers indicate number of neuronal cultures (N=2) and total number of cells analyzed (n=24, n=20); axonal length: Mann-Whitney U = 32, *p*<0.0001, Mann-Whitney test; axonal branches: Mann-Whitney U = 53, *p*<0.0001, Mann-Whitney test. **(B)** Overview coronal sections in grey scale stained with anti-RFP antibody of an empty vector and a RHEBp.P37L mouse brain *in utero* electroporated on the left S1 and magnification of the axon terminals on the contralateral S1; scale bars: 500 μm. **(C)** Representative images of ipsilateral and contralateral S1 area of an empty vector and a RHEBp.P37L mouse coronal section (P50) with quantification of the axonal projections across the different layers in the contralateral cortex measured as normalized fluorescent intensity of the tdTomato signal; numbers in the legend indicate number of targeted mice (N=3, N=4) and number of contralateral pictures (n=10, n=23) analyzed; data are presented as mean (thick line) ± SEM (shading area); interaction group condition/cortical layers: F(9, 279)=13.96, *p*<0.0001, mixed-effects analysis; control *vs* RHEBp.P37L L2/3 (bin2-3 from the top): *p*<0.0001; control *vs* RHEBp.P37L L5-L6: bin7, *p*=0.0074, bin8, *p*<0.0001, bin 9, *p*=0.002; Bonferroni multiple comparisons test. * *p*<0.01, **** *p*<0.0001; scale bars: 50 μm **(A)**, 500 μm **(B)**, 100 μm **(C)**.

*In vivo*, axons from callosal projection neurons originating from the superficial layers of the SScx project to the homotopic contralateral hemisphere, where they mostly innervate L2/3 and L5 pyramidal neurons (Fenlon et al., 2017; Petreanu et al., 2007). They also send collaterals to L2/3 and, more strongly, L5 pyramidal neurons within the same column ipsilaterally, participating in local circuitry (Fame et al., 2011; Petreanu et al., 2007). Therefore, it is conceivable that *in vivo* overexpression of RHEBp.P37L affects callosal projections to the non-targeted contralateral hemisphere. Analysis of the contralateral callosal axonal growth in matched coronal sections with comparable targeting revealed that upon RHEBp.P37L overexpression, axonal terminals in the contralateral hemisphere, show a broader distribution compared to controls, reaching the primary (S1) and secondary (S2) SScx (Figure 5B). Furthermore, a significant difference was found in the distribution of the axonal terminals across the different layers in the contralateral hemisphere. While in the control condition most of the terminals were located in L2/3, with a lower abundance in L5 (Fenlon et al., 2017), in the RHEBp.P37L mice we found that most of the terminals were located in the deeper layers of the cortex, suggesting an improper cortical connectivity (Figure 5C). Furthermore, zooming in on the axonal projections on the contralateral cortex revealed the presence of enlarged *boutons* and terminals in the RHEBp.P37L mice that were both Synapsin-1 and VGLUT1 positive, markers for synaptic vesicles and glutamatergic neurons, respectively (**Figure 5-figure supplement 1**).

To investigate if the contralateral axonal projections with these synaptic terminals showing altered morphology are functional, we made use of optogenetics. We used IUE to introduce channelrhodopsin-2 (pCAGGS-ChR2-Venus) (Petreanu et al., 2007) together with either the empty vector control or the RHEBp.P37L construct in targeted neurons and recorded the postsynaptic responses (EPSCs) to widefield optogenetic stimulation by patch-clamping L2/3 and L5 pyramidal neurons in the (non-targeted) contralateral S1 where axonal terminals could be observed (Figure 6A **and Figure 6-figure supplement 1A**). Analyzing the amplitude of EPSCs following optogenetic stimulation in L5 and L2/3 of the contralateral S1, we observed an overall increase in response in the RHEBp.P37L condition compared to the empty vector control condition (see **Supplementary table 3** for statistics) (Figure 6B). When analyzing the total charge of the compound postsynaptic response we observed similar response patterns (**Supplementary table 3** for statistics) (Figure 6B). Bath application of tetrodotoxin (TTX) in the RHEBp.P37L group decreased the post-synaptic responses evoked by photo-stimulating ChR2 expressing fibers to noise level, which is indicative of action potential driven neurotransmitter release (**Figure 6-figure supplement 1B**). The basic properties (resting membrane potential [Vm] and membrane resistance [Rm]) of L2/3 and L5 contralateral cells in empty vector control and RHEBp.P37L conditions were not different (**Supplementary table 3** for statistics). These data suggest increased synaptic connectivity to the contralateral S1 upon overexpression of RHEBp.P37L.

**Figure 6.**
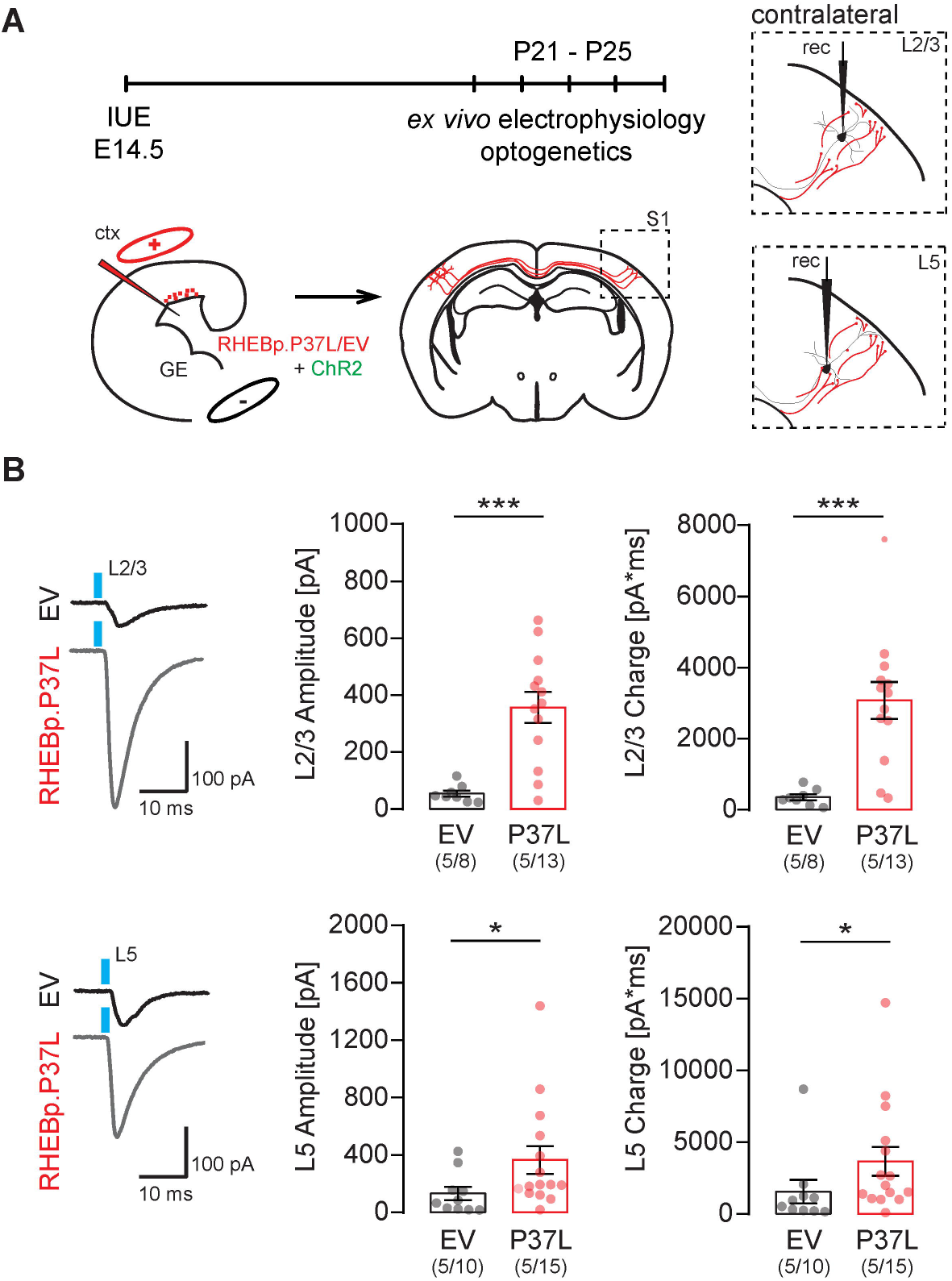
Overexpressing RHEBp.P37L increases synaptic connectivity on the contralateral hemisphere. **(A)** Schematic representation of the timeline and experimental conditions of the IUE and *ex vivo* whole-cell patch clamp recordings in contralateral L2/3 and L5 upon wide-field optogenetic stimulation. **(B)** Example traces and analysis of the compound postsynaptic responses after photostimulation (blue light), showing the postsynaptic response amplitudes and total charge in contralateral L2/3 and L5 in empty vector (EV) and RHEBp.P37L expressing slices; numbers in the graph indicate number of targeted mice (N=5) and number of cells (n=8, n=13, n=10, n=15) analyzed; data are presented as mean ± SEM and single data points indicate the values of each cell; for statistics see **Supplementary table 3**; * *p*<0.05, *** *p*<0.001.

### Loss of axonal projections or blocking vesicle release of RHEBp.P37L expressing neurons is sufficient to stop seizures

Having shown that the RHEBp.P37L expressing neurons show stronger axonal innervation and synaptic connectivity to neurons in the contralateral hemisphere, we investigated whether these altered neuronal projections drive the seizures. To assess this, we made use of the Tetanus toxin light chain, known to specifically cleave the SNARE-complex protein Synaptobrevin/VAMP2 (Syb2) (Schiavo et al., 1992). VAMP2 is part of the SNARE complex that allows synaptic vesicles fusion and the release of neurotransmitters (Gaisano et al., 1994) and recently it has been shown to mediate the vesicular release of Brain Derived Neurotrophic Factor (BDNF) from axon and dendrites, thereby regulating proper cortical connectivity (Shimojo et al., 2015). Intrinsic neuronal activity during early brain development is crucial for axonal growth and branching, and blocking synaptic transmission using Tetanus toxin interferes with proper cortical axonal formation, resulting in the reduction and disappearance of axonal projections (Wang et al., 2007). Indeed, when RHEBp.P37L was co-transfected with a Tetanus toxin construct (TeTxLC) that is active during embryonic development, we observed a complete block of callosal axonal growth (Figure 7A, 7B). Furthermore, the mice targeted with the RHEBp.P37L and TeTxLC constructs did not develop any seizures, suggesting that the abnormal axonal connectivity might mediate the expression of seizures in our mouse model (Figure 7B).

**Figure 7.**
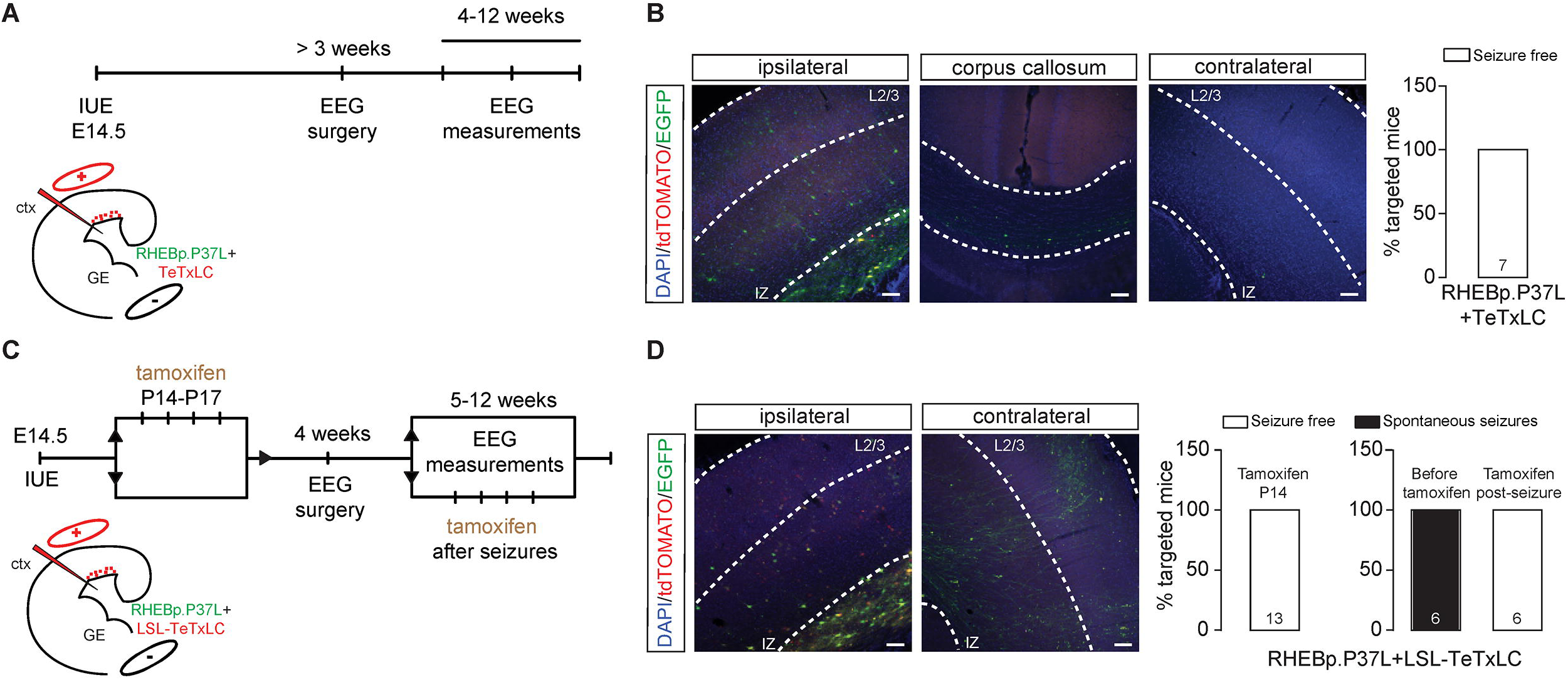
Loss of axonal projections or blocking vesicle release of RHEBp.P37L expressing neurons is sufficient to stop seizures. **(A) (C)** Schematic representation of the timeline of the IUE, EEG surgery and measurements and timepoints of tamoxifen administration in **(C)**. **(B)** Example figures of ipsilateral targeted S1, corpus callosum and contralateral S1 of an adult mouse (12 weeks) *in utero* electroporated with the RHEBp.P37L (construct expressing EGFP, in green) and a Tetanus toxin construct (TeTxLC, construct expressing tdTomato, in red). Note the absence of axonal projections on the contralateral side. The bar graph shows percentage of seizure-free targeted mice measured with EEG until 12 weeks of age. Numbers in the bar graph indicate number of mice. **(D)** Example figures of ipsilateral targeted area (left) and contralateral cortex of an adult mouse (12 weeks) *in utero* electroporated with the RHEBp.P37L (in green) and a LSL-Tetanus toxin construct (LSL-TeTxLC, in red) and injected with tamoxifen starting at P14. The bar graph shows percentage of targeted mice developing seizures upon early tamoxifen injection (P14) or upon tamoxifen administration after detecting seizures and measured with EEG until 12 weeks of age (see **Supplementary table 4** for details on the timeline of the experiment). Numbers in the bar graphs indicate number of mice. Scale bars: 100 μm.

The complete loss of callosal axonal branching upon embryonic activation of TeTxLC, prevented us from testing whether increased synaptic transmission drives seizure development. Therefore, to enable activation of the Tetanus toxin upon Tamoxifen injection at post-developmental stages, we generated an inducible LSL-TeTxLC construct (Figure 7C) and co-transfected this construct with RHEBp.P37L and the CAGG-ERT2CreERT2 vector (see Figure 4E). This allowed us to assess whether, once (abnormal) axonal projections are established, blocking vesicular release either prevents the development of seizures, or stops seizures once they have developed. Activation of the Tetanus toxin during postnatal development, but before seizure onset (P14), completely prevented the development of seizures while allowing the axons to grow and branch to the contralateral side (Figure 7D). Administering tamoxifen in 5 weeks old mice, when the cortical connectivity is complete and after the mice showed seizures revealed that epilepsy is not an irreversible process (Figure 7D). Already after 2 days of tamoxifen administration, 3 out of 6 mice (**Supplementary table 4**) stopped showing any seizures and 2 weeks after the last tamoxifen injection all mice appeared to be seizure free (**Supplementary table 4**). These results indicate that inhibiting synaptic transmission by blocking vesicular release from the targeted cells is enough to stop the occurrence of seizures in our mouse model.

### Neurons in the contralateral homotopic cortical area in RHEBp.P37L expressing mice show increased excitability

To obtain more insight into the cellular mechanisms that underlie epilepsy in our model, we used whole-cell patch clamp to measure intrinsic physiological properties of the RHEBp.P37L expressing neurons, of (ipsilateral) neurons directly surrounding the targeted cells, and of the contralateral neurons in homotopic cortical areas (Figure 8A). Whole cell patch clamp recordings were performed by recording from pyramidal neurons in S1 of 3 weeks old mice. For the RHEBp.P37L expressing neurons (tdTomato positive), we recorded only from neurons that managed to migrate out to L2/3 of S1 to be able to compare their physiological properties with ‘empty vector’ control cells in L2/3 that expressed the tdTomato gene without expressing the RHEBp.P37L protein (Figure 8A). RHEBp.P37L expressing neurons showed an increase in the capacitance (*Cm*) compared to empty vector control cells (Figure 8B and see **Supplementary table 5** for statistics), which is consistent with the increase in soma size (median of control empty vector cells L2/3: 1.005, n cells=22; median RHEBp.P37L cells L2/3: 1.377, n cells=24; Mann-Whitney U = 105, *p*=0.0003, Two-tailed Mann-Whitney test, data not shown). Additionally, the membrane resistance (*Rm*) was decreased, whereas the resting membrane potential (*Vm*) was unchanged compared to empty vector control cells (Figure 8B and see **Supplementary table 5** for statistics). Depolarizing the neurons with increasing current injections, showed that the excitability of cells expressing the empty vector were not different from non-targeted neurons in the same mice or compared to non-targeted mice (**Figure 8-figure supplement 1**). In contrast, RHEBp.P37L expressing neurons were hypoexcitable compared to control neurons measured in mice expressing the empty vector as well as to non-targeted neurons ipsilateral and contralateral (Figure 8C and see **Supplementary table 5** for statistics), without a change in the threshold *Vm* to fire action potentials (F (3, 94) = 0.59, *p*=0.62, non-significant, One-way ANOVA). This result is again in agreement with the observed increased soma size and concomitant increased cell capacitance and decreased membrane resistance. Notably, while ipsilateral non-transfected neurons surrounding the RHEBp.P37L expressing neurons in mice did not show changes in excitability compared to empty vector control, non-transfected neurons in L2/3 on the contralateral hemisphere showed a significant increase in excitability (Figure 8C and see **Supplementary table 5** for statistics), suggesting that the ectopic cells affect long-range connected neurons.

**Figure 8.**
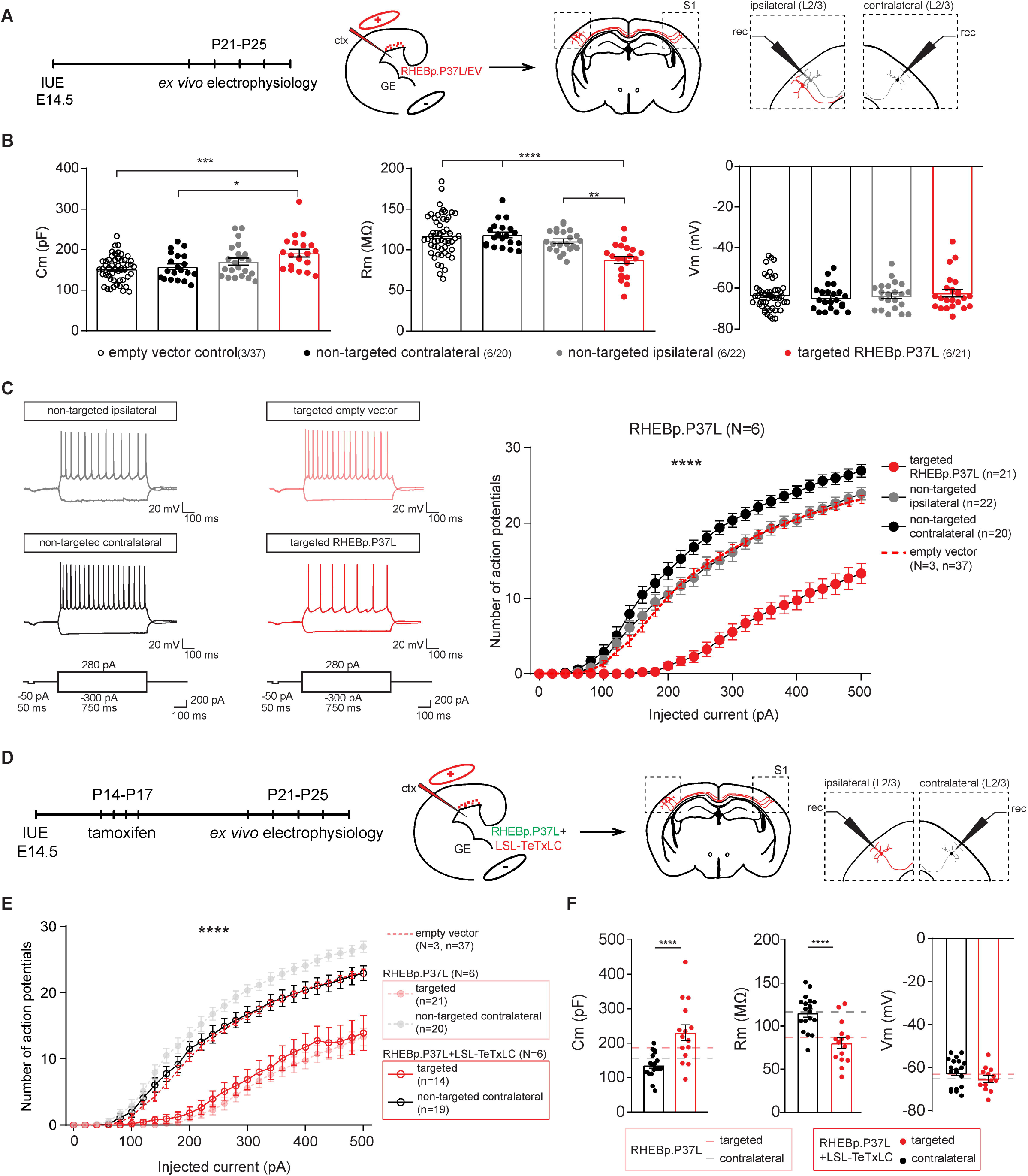
Neurons in the contralateral homotopic cortical area in RHEBp.P37L mice show increased excitability that can be reversed by blocking vesicles release. **(A)** Schematic representation of the timeline and experimental conditions of the IUE and *ex vivo* whole-cell patch clamp recordings showing the targeted cells patched in the targeted S1, L2/3 and non-targeted L2/3 cells in the ipsilateral and contralateral sides. **(B)** Analysis of the passive membrane properties (capacitance [Cm], membrane resistance [Rm] and resting membrane potential [Vm]) of pyramidal cells in L2/3 (targeted and non-targeted) of control empty vector mice and targeted and non-targeted pyramidal cells in L2/3 of RHEBp.P37L mice; numbers in the legend indicate number of targeted mice (N=3, N=6) and number of cells (n=37, n=20, n=22, n=21) analyzed; data are presented as mean ± SEM and single data points indicate the values of each cell; for statistics see **Supplementary table 5**). **(C)** Example traces and number of action potentials in response to increasing depolarizing currents; number of mice and cells is as indicated in **(B)**; data are presented as mean ± SEM and the red dashed line represents the pooled mean value ± SEM of targeted and non-targeted cells in empty vector control mice (N=3) shown separately in **Figure 8-figure supplement 1**, for comparison; for statistics see **Supplementary table 5**). **(D)** Schematic representation of the timeline and experimental conditions IUE, tamoxifen injections and *ex vivo* whole-cell patch clamp recordings in L2/3 of ipsilateral and contralateral S1 cortex. **(E)** Number of action potentials in response to increasing depolarizing currents of cells expressing both RHEBp.P37L and LSL-TeTxLC in L2/3 ipsilateral S1 and non-targeted cells in L2/3 contralateral S1; data are presented as mean ± SEM and dashed lines represent the mean values ± SEM of the pooled control cells from empty vector mice shown in **Figure 8**-**figure supplement 1** and of the RHEBp.P37L mice from Figure 8C, for comparison; N=number of mice, n=number of cells analyzed; for statistics see **Supplementary table 6**. **(F)** Analysis of passive membrane properties (Cm, Rm and Vm) of pyramidal cells in L2/3 of mice targeted with RHEBp.P37L and LSL-TeTxLC in ipsilateral S1 and non-targeted cells on the contralateral side; data are presented as mean ± SEM and the dashed lines indicate the mean values of capacitance, membrane resistance and resting membrane potential of RHEBp.P37L targeted cells in L2/3 and contralateral cells shown in Figure 8B, for comparison; for statistics see **Supplementary table 6**. * *p*<0.05, ** *p*<0.01, *** *p*<0.001, **** *p*<0.0001.

To experimentally address if the aberrant connectivity could cause the increase in excitability in neurons on the contralateral cortex, we again made use of the inducible LSL-TeTxLC construct and co-transfected this construct with RHEBp.P37L and the CAGG-ERT2CreERT2 vector to enable activation of the Tetanus toxin upon Tamoxifen injection at post-developmental stages (See Figure 7C). Whole-cell patch clamp recordings revealed that activating the Tetanus toxin early during development (P14) (Figure 8D), completely reversed the hyperexcitability phenotype of the contralateral non-targeted cells observed in the RHEBp.P37L mice (Figure 8E) while the targeted cells co-transfected with the RHEBp.P37L and the Tetanus toxin maintained the hypoexcitable phenotype and the basic properties observed in the RHEBp.P37L group (Figure 8E-F and see **Supplementary table 6** for statistics). Taken together, these data indicate that the abnormal axonal connectivity caused by RHEBp.P37L overexpression is the primary driver of the hyperexcitability phenotype of contralateral L2/3 pyramidal neurons, which in turn could be the main driver of epilepsy.

## DISCUSSION

In this study, we investigated the mechanisms behind the spontaneous tonic-clonic seizures in a mouse model that was generated by spatially and temporally restricted overexpression of a mTOR-related ID mutation in *RHEB* (Reijnders et al., 2017). We showed that the RHEBp.P37L mutant is resistant to inhibition by the TSC complex, and that restricted overexpression causes mTORC1 hyperactivity and the development of heterotopia with typical cellular features of human MCD such as enlarged dysplastic neurons with altered morphology and mTORC1 activation. Furthermore, the presence of cortical malformations is accompanied by the development of spontaneous tonic-clonic seizures and alterations of the cortical brain dynamics that are rescued by administration of rapamycin, an mTOR inhibitor. Using a pharmacological and genetic approach we showed that the presence of the cortical malformation by itself is neither necessary nor sufficient to induce epilepsy, while blocking either mTOR activity or vesicle release from these cells is enough to stop or prevent seizures.

Similar to previously generated IUE mouse models of MCD, our model developed clear heterotopia, strikingly resembling focal human cortical malformations, associated with mTORC1 hyperactivity and reliable spontaneous seizures (Hanai et al., 2017; Hsieh et al., 2016; Lim et al., 2015; Park et al., 2018; Ribierre et al., 2018). The malformation in our mouse model is characterized by white matter heterotopia and neuronal misplacement across the different cortical layers, but maintains the molecular fingerprint belonging to L2/3 neurons. However, it is difficult to categorize it as a specific type of MCD because it expresses characteristics of both FCD type I and type IIa (with no Balloon cells observed) (Barkovich et al., 2012). Nonetheless, the targeted cells have features common to several types of mTOR dependent MCD, including enlarged and dysplastic cells with mTORC1 hyperactivation (Crino, 2011).

Previously it has been shown that brain wide activation of the mTOR pathway is sufficient to induce seizures in the absence of any cortical malformations (Abs et al., 2013). However, these models do not address the role of mTOR signaling in MCD related pathophysiology. To address that, an elegant IUE mouse model was generated which expressed the constitutive active RHEBp.S16H protein. These mice showed a migration deficit resembling FCD and spontaneous epilepsy (Hsieh et al., 2016). Using this mouse model, it was also shown that the presence of a cortical malformation is not necessary to induce seizures (Hsieh et al., 2016). Notably, these mice did not show epilepsy when the SScx was targeted, and hence the investigators suggested that the SScx might be a non-epileptogenic area. This is in contrast with our mouse model using the human ID-related RHEBp.P37L mutant, where targeting the S1 area of the SScx, reliably induces seizures. One reason for these contradictory findings might be the effect of the mutation on RHEB function, as we showed that the RHEBp.S16H mutant could be partially inhibited by the TSC complex, whereas the RHEBp.P37L mutant was insensitive to TSC regulation.

Our mouse model offers a good tool to test novel AEDs *in vivo*. However, considering the variability in the number of seizures exhibited, it will be beneficial to focus on different parameters when assessing the potential therapeutic efficiency of AEDs. For this purpose, the *theta* frequency oscillation, which we found to be affected and normalized upon rapamycin treatment, represents a good biomarker for assessing the potential therapeutic value of treatments in our mouse model (Chauvière et al., 2009; Milikovsky et al., 2017).

Everolimus and rapamycin (Sirolimus) have been shown in randomized controlled trials to be beneficial for treating TSC associated epilepsy (Iris E Overwater et al., 2019; Overwater et al., 2016), but not for treating the cognitive deficits (Krueger et al., 2017; Iris E. Overwater et al., 2019). In this study we investigated the potential of a short prenatal rapamycin treatment in improving both malformation defects and epilepsy, but preventing the possible side effects (developmental delays and poor gain weight) (Tsai et al., 2013). We showed that a 2-day rapamycin treatment during a critical time point of prenatal development can cause a substantial improvement of the cortical malformation defects and prevent the development of seizures in almost 50% of the cases. Future studies will have to assess if a combination of prenatal and postnatal treatment with rapamycin in mice can be sufficient to significantly reduce the epileptic events, as shown for brain malformations, without causing major side effects (Tsai et al., 2013; Way et al., 2012).

Surgery is often an alternative to AEDs for treating MCD-related epilepsy. Human electrophysiological findings show that seizures can often have multiple starting points, besides the brain lesion itself (Chassoux et al., 2008; Major et al., 2009). Therefore, from a clinical point of view, it is important to determine whether seizures originate from cells surrounding the cortical malformation. Even though EEG and LFP do not have the spatial resolution to assess the primary epileptogenic zone in our model, we showed that persistent mTORC1 hyperactivation in the targeted cells is the primary cause of epilepsy. In fact, genetically removing the RHEBp.P37L mutant, either before or after seizure development, was sufficient to prevent or stop the epilepsy.

Surprisingly, when exploring the causes of epileptogenesis, we observed that the neurons expressing the RHEBp.P37L are hypoexcitable, which is consistent with the increase in soma size but does not provide an obvious physiological explanation for the seizures observed in our mouse model. However, we observed a clear increase of intrinsic excitability and in postsynaptic responses upon optogenetic stimulation of RHEBp.P37L cells in contralateral homotopic S1 cells. This suggests that RHEBp.P37L expressing cells induce cellular changes in anatomically connected neurons, which might underlie, or at least exacerbate, the epilepsy phenotype. Notably, these alterations extend well beyond the cells surrounding the cortical malformation, as we found physiological changes were present contralateral to the targeted side. Considering the abnormal axonal connectivity seen in our mouse model, this raises the possibility that other anatomically connected cortical and sub-cortical areas not analyzed in this study might also be affected, thereby providing an explanation for how a small percentage of targeted hypoexcitable cells, independent of their location, can lead to generalized epilepsy. Therefore we propose a model in which subtle microscopic alterations and aberrant connectivity, either through an increase in synaptic connections or an increase in the strength of synaptic contacts caused by mTOR hyperactivity, are sufficient to drive epileptogenesis.

By increasing axonal connectivity, RHEBp.P37L expressing neurons could potentially alter synaptically connected neurons through neurotransmitter release. But they can also affect neighboring (including synaptically non-connected) cells through the release of extracellular vesicles such as exosomes (Budnik et al., 2016). The vesicles might mediate pathogenicity as was previously shown *in vitro* (Patel et al., 2015). With the use of Tetanus toxin, we showed that the effects on the contralateral side are directly driven by the abnormal enhanced axonal connectivity, since blocking vesicle release specifically from the RHEBp.P37L expressing neurons, completely rescued the epilepsy and normalized the intrinsic firing properties of the non-targeted contralateral neurons. Tetanus toxin is primarily used to block synaptic transmission due to its effect on neurotransmitter release, acting on the SNARE complex protein VAMP2 (Schiavo et al., 1992). Given the observed increased axonal connectivity and the finding that distally connected cells were physiologically affected, this strongly suggest that neurotransmitter mediated communication is primarily causing the epilepsy phenotype. This notion is further supported by the optogenetics experiments that showed increase postsynaptic responses upon stimulating the RHEBp.P37L expressing neurons. While it has been proposed that specific tetanus insensitive VAMP proteins (such as VAMP7) are involved in the release of exosomes into the extracellular space (Fader et al., 2009), we cannot exclude the additional contribution of other types of vesicles to the observed phenotype. Recently it was shown that Tetanus toxin sensitive SNAREs also drive the release of BDNF (Shimojo et al., 2015). Some studies suggest that BDNF might contribute to epileptogenesis (Binder et al., 2001), suggesting that abnormal BDNF signaling could further increase the epileptic phenotype seen in our mouse model. Understanding the contribution of these different signaling pathways is important for the development of targeted therapeutic strategies to treat MCD associated epilepsy.

In summary, in this study we show that restricted overexpression of a hyperactive RHEB mutant that was previously identified in patients with ID, megalencephaly and epilepsy, strongly mimics the human MCD-like phenotype with mTOR pathway hyperactivity and seizures. We provided pharmacological and genetic evidence that the cortical malformation *per se* is neither necessary nor sufficient to induce seizures. Furthermore, we show that only a few neurons with increased mTOR activity can be the driving force behind MCD-related epilepsy through aberrant connectivity, resulting in increased excitability of distant non-targeted neurons, which can be reversed by blocking vesicular release.

## MATERIALS AND METHODS

### Mice

Unless subjected to a surgical procedure, all experimental mice were kept group-housed in IVC cages (Sealsafe 1145T, Tecniplast) with bedding material (Lignocel BK 8/15 from Rettenmayer) on a 12/12 hr light/dark cycle at 21°C (±1°C), humidity at 40-70% and with food pellets (801727CRM(P) from Special Dietary Service) and water available *ad libitum*. For the neuronal cultures, FvB/NHsD females were crossed with FvB/NHsD males (both ordered at 8-10 weeks old from Envigo). For the IUE, females FvB/NHsD (Envigo) were crossed with males C57Bl6/J (ordered at 8-10 weeks old from Charles River). Both females and males from the *in utero* electroporation litters were included in the experiments and no prescreening for successful electroporation was performed before recruitment in the studies. Young (starting from P7) and adult mice were used and the specific age for each experiment is indicated either in the results section or in the figures’ legends. Activation of the ERT2CreERT2 fusion protein (Matsuda and Cepko, 2007) was achieved by intraperitoneal administration of tamoxifen for 4 consecutive days (0.1 mg/g of bodyweight) dissolved in sunflower oil (20 mg/ml) at the ages specified in the results section and in the figures. For inhibition of the mTOR pathway, rapamycin (Sigma-Aldrich) was dissolved in dimethylsulfoxide (10 mg/ml) and injected intraperitoneally in adult mice (> 4 weeks) for postnatal experiments (10 mg/kg) or subcutaneously in pregnant females (E15.5/E16.5) for prenatal experiments (1 mg/kg).

All animal experiments were conducted in accordance with the European Commission Council Directive 2010/63/EU (CCD approval AVD1010020172684).

### HEK293T cell cultures and transfection

HEK293T cells were grown in Dulbecco’s modified Eagle medium (DMEM; Lonza, Verviers, Belgium) supplemented with 10% fetal bovine serum, 50 U/ml penicillin and 50 μg/ml streptomycin in a 5% CO_2_ humidified incubator at 37°. Before transfection, 1 x 10^5^ HEK293T cells were seeded per well of 6-well culture dishes and transfected 24 hours later with expression constructs encoding the *RHEB* variants (0.2 μg), the S6K reporter (0.2 μg), *TSC1* (0.2 μg) and *TSC2* (0.2 μg) using Lipofectamine 2000 (Invitrogen, Carlsbad, CA, USA). To ensure that a total of 0.8 μg plasmid DNA was added per well, empty pcDNA3 vector was included where necessary. The day after transfection, the growth medium was replaced with DMEM without glucose and incubated for a further 4 hours prior to harvesting and western blot analysis.

### Western blotting

After transfection, HEK293T cells were transferred on ice, washed with PBS (4 °C) and lysed in 70 μl 50 mM Tris-HCl pH 7.6, 100 mM NaCl, 50 mM NaF, 1% Triton X100 in the presence of protease and phosphatase inhibitors (Complete, Roche Molecular Biochemicals, Woerden, The Netherlands). Cell lysates were subjected to immunoblotting using the following primary antibodies: anti-RHEB mouse monoclonal (Groenewoud et al., 2013), anti-TSC1 and TSC2 rabbit polyclonal (Van Slegtenhorst et al., 1998), T389-phosphorylated S6K (1A5, #9206, Cell Signaling Technology), and rabbit anti-myc (#2272, Cell Signaling Technology), all 1:1000. Primary antibody binding was assessed by incubation with goat anti-rabbit (680 nm) and anti-mouse (800 nm) conjugates (1:15000, Li-Cor Biosciences, Lincoln, USA) followed by detection on an Odyssey near-infrared scanner (Li-Cor Biosciences).

### Neuronal primary hippocampal cultures and transfection

Primary hippocampal neuronal cultures were prepared from FvB/NHsD wild type mice according to the procedure described in (Banker and Goslin, 1988). Neurons were transfected at 1 day *in vitro* (DIV1) with the following DNA constructs: control empty vector (1.8 μg per coverslip) and RHEB p.P37L (2.5 μg per coverslip). Plasmids were transfected using Lipofectamine 2000 according to the manufacturer’s instructions (Invitrogen).

### Plasmids

cDNA encoding the *RHEB* (NM_005614.3) c.110C>T (p.P37L) mutation was synthesized by GeneCust. The c.46-47CA>TG (p.S16H) variant was generated by site-directed mutagenesis (Invitrogen) using the following primers: Fw 5’ – gcgatcctgggctaccggCAtgtggggaaatcctcatt – 3’ and Rev 5’ – aatgaggatttccccacaTGccggtagcccaggatcgc – 3’. All *RHEB* gene variants were cloned in our dual promoter expression vector using AscI and PacI restriction sites (Reijnders et al., 2017) and the empty vector used as control refers to the dual promoter expression vector without a gene inserted and expressing either tdTOMATO or EGFP (specified in the figures or in the figures’ legends). Expression constructs for TSC1, TSC2 and a myc-tagged S6K reporter were as described previously (Dufner Almeida et al., 2019). The following DNA plasmids were obtained from Addgene: pGEMTEZ-TeTxLC (Addgene plasmid #32640; http://n2t.net/addgene:32640; RRID:Addgene_32640) (Yu et al., 2004); RV-CAG-DIO-EGFP (Addgene plasmid #87662; http://n2t.net/addgene:87662; RRID:Addgene_87662) (Ciceri et al., 2013); pCAG-ERT2CreERT2 (Addgene plasmid #13777; http://n2t.net/addgene:13777; RRID:Addgene_13777) (Matsuda and Cepko, 2007); pCAGGS-ChR2-Venus (Addgene plasmid #15753; http://n2t.net/addgene:15753; RRID:Addgene_15753) (Petreanu et al., 2007). The TeTxLC was isolated by PCR using the following primers: Fw 5’ – taagcaggcgcgccaccatgccgatcaccatcaacaa – 3’ and Rev 5’ – gccatggcggccgcgggaattcgat – 3’ and inserted in our dual promoter expression vector using AscI and NotI restriction sites. To generate the loxP-STOP-loxP (LSL) constructs (loxP-STOP-loxP-*RHEB* p.P37L and loxP-STOP-loxP-TeTxLC) the LSL sequence was obtained from the Ai6 CAG-Floxed ZsGreen in Rosa 26 targeting vector (Addgene plasmid #22798; http://n2t.net/addgene:22798; RRID:Addgene_22798) using multiple cloning sites and inserted just after the CAGG promoter and before the beginning of the gene in our dual promoter expression vector containing either *RHEB*p.P37L or TeTxLC. The floxed *RHEB* p.P37L construct was generated by introducing two loxP site sequences before the CAGG promoter and at the end of the *RHEB*p.P37L gene, with the same orientation to ensure proper deletion. To achieve this, the following couples of oligonucleotides were used for annealing: Fw 5’-cgcgtATAACTTCGTATAGCATACATTATACGAAGTTATg −3’, Rev: 5’-ctagcATAACTTCGTATAATGTATGCTATACGAAGTTATa −3’; Fw: 5’-taaATAACTTCGTATAGCATACATTATACGAAGTTATg −3’, Rev: 5’-tcgacATAACTTCGTATAATGTATGCTATACGAAGTTATttaat −3’.

### In utero electroporation

IUE was performed as described previously (Saito and Nakatsuji, 2001). Pregnant FvB/NHsD mice at E14.5 of gestation were used to target the progenitor cells giving rise to pyramidal cells of the layer 2/3. Each *RHEB* DNA construct (including the LSL and floxed conditions) was diluted to a final concentration of 0.5 μg/μl in fast green (0.05%), while other plasmids were diluted to a concentration of 1.5-2 μg/μl. The DNA solution was injected into the lateral ventricle of the embryos while still *in utero*, using a glass pipette controlled by a Picospritzer ® III device. When multiple constructs were injected, a mixture of plasmids was prepared to achieve a final concentration of 1.5-2 μg/μl, keeping the *RHEB* concentration constant throughout all the experiments. To ensure proper electroporation of the injected constructs (1-2 μl) into the progenitor cells, five electrical square pulses of 45V with a duration of 50 ms per pulse and 150 ms inter-pulse interval were delivered using tweezer-type electrodes connected to a pulse generator (ECM 830, BTX Harvard Apparatus). The positive pole was placed to target the developing somatosensory cortex. Animals of both sexes were used to monitor seizure development, for *ex vivo* electrophysiology experiments, or for histological processing with no exclusion criteria determined by a postnatal screening of the targeting area.

### Immunostainings

For immunocytochemistry analysis, neuronal cultures were fixed 3 days post-transfection with 4% paraformaldehyde (PFA)/4% sucrose, washed in PBS and incubated overnight at 4°C with primary antibodies in GDB buffer (0.2% BSA, 0.8 M NaCl, 0.5% Triton X-100, 30mM phosphate buffer, pH7.4). Mouse pan anti-SMI312 (1:200, BioLegend, #837904) was used to stain for the axon and, after several washings in PBS, donkey anti-mouse-Alexa488 conjugated was used as secondary antibody diluted in GDB buffer for 1 hour at room temperature (1:200, Jackson ImmunoResearch). Slides were mounted using mowiol-DABCO mounting medium.

For the staining of brain tissue sections, mice were deeply anesthetized with an overdose of Nembutal and transcardially perfused with 4% PFA in PB. Brains were extracted and post-fixed for 1 hour in 4% PFA. They were then embedded in gelatin and cryoprotected in 30% sucrose in 0.1 M Phosphate Buffer (PB) overnight, frozen on dry ice, and sectioned using a freezing microtome (40 μm thick). Immunofluorescence was performed on free-floating sections that were first washed multiple times in PBS and blocked in 10% normal horse serum (NHS) and 0.5% Triton X-100 in PBS for 1 hour at room temperature. Primary antibodies diluted in PBS containing 2% NHS and 0.5% Triton X-100 were added at room temperature overnight. The day after, sections were washed three times with PBS and secondary antibodies were added diluted in PBS containing 2% NHS and 0.5% Triton X-100. After washing in PBS and 0.05 M PB, sections were counterstained with 4′,6-diamidino-2-phenylindole solution (DAPI, 1:10000, Invitrogen) before being washed in PB 0.05 M and mounted on slides using chromium (3) potassium sulfate dodecahydrate (Sigma-Aldrich) and left to dry. Finally, sections were mounted on glass with mowiol (Sigma-Aldrich).

Biocytin labelling was achieved by fixating the patched slices overnight in 4% PFA in PB at 4°. Slices were then washed multiple times in PBS and incubated with Alexa488-Streptavidin (1:200; #016-540-084, Jackson ImmunoResearch) or AlexaCy5-Streptavidin (1:200; #016-170-084, Jackson ImmunoResearch) overnight at 4°. The next day, after washing in PBS and 0.05 M PB, sections were counterstained with DAPI (1:10000, Invitrogen) and mounted on glass with mowiol (Sigma-Aldrich).

When performing Nissl stainings, few selected free floating sections corresponding to the Somatosensory cortex were mounted on glass using chromium (3) potassium sulfate dodecahydrate (Sigma-Aldrich) and left to dry overnight. Slides were stained in 0.1 % Cresyl Violet for 4-10 minutes, then rinsed briefly in tap water to remove excess stain, dehydrated in increasing percentages of alcohol, cleared with xylene and covered using Permount (Fisher Scientific).

The primary antibodies used in this study to stain for the specific targets indicated for each experiment in the figures’ legends were: anti-rabbit pS6 (Ser 240/244), 1:1000; Cell signaling, catalog #5634; anti-rabbit RFP, 1:2000; Rockland, catalog 600-401-379; anti-rabbit RHEB, 1:1000, Proteintech Group Inc., catalog 15924-1-AP; anti-rabbit CUX1, 1:1000; Proteintech Group Inc., catalog 11733-1-AP; anti-rat CTIP2, 1:200; Abcam, catalog ab18465; anti-rabbit NeuN, 1:2000; Millipore catalog ABN78 (RRID: AB_10807945); anti-mouse SATB2, 1:1000; Santa cruz, catalog sc-81376; anti-rabbit synapsin 1, 1:1000; Merck Millipore, catalog #AB1543P; anti-guinea pig VGLUT1, 1:1000; Merck Millipore, catalog #AB5905; Secondary antibodies used were: donkey anti rabbit 488, catalog #711-545-152; donkey anti rabbit 647, catalog #711-605-152; donkey anti rabbit Cy3, catalog #711-165-152; donkey anti mouse 488, catalog #715-545-150; donkey anti mouse 647, catalog #715-605-150; donkey anti rat Cy5, catalog #712-175-150; donkey anti guinea pig 647, catalog #706-605-148; all from Jackson ImmunoResearch, 1:200.

### LFP and EEG recordings

Starting from 3 weeks of age surgeries were performed according to the procedures described in (Koene et al., 2019; Kool et al., 2019). After at least three days of recovery from the EEG surgical procedure, mice were connected to a wireless EEG recorder (NewBehavior, Zurich, Switzerland) for 24 hours per day for at least two consecutive days (one session of recordings). EEG recordings were manually assessed by two different researchers blind for the genotypes to check for occurrence of seizures, defined as a pattern of repetitive spike discharges followed by a progressive evolution in spike amplitude with a distinct post-ictal depression phase, based on the criteria described in (Kane et al., 2017). If no seizures were detected during the first week *post*-surgery, mice were recorded for another session of 48-56 hr for a maximum of four sessions over four weeks *post*-surgery. During the days in which no EEG recordings were performed, mice were monitored daily to assess for the presence of behavioural seizures and discomfort.

For the LFP recordings, two days after the surgical procedure, mice were head-fixed to a brass bar suspended over a cylindrical treadmill to allow anaesthesia-free recording sessions and placed in a light-isolated Faraday cage as described in (Kool et al., 2019). Mice were allowed to habituate to the set-up before proceeding to the recording. LFP measurements were acquired every day in sessions of 20-30’ for five or eight consecutive days, using the Open Ephys platform with a sampling rate of 3 kS/s and a band pass filter between 0.1 and 200 Hz. For the power spectrum analysis, the average power density spectrum of all the days of recording was obtained using MATLAB software (MathWorks; RRID:SCR_001622). The mean relative power was calculated over four frequency bands relative to the total power: delta (2–4 Hz), theta (4–8 Hz), beta (13–30 Hz), and gamma (30–50 Hz).

At the end of each experiment, mice were sacrificed for immunohistological analysis to assess electrodes’ positioning, amount of targeting and efficiency of cre-dependent recombination when tamoxifen was administered.

### *Ex vivo* slice electrophysiology for excitability

P21-P25 mice of both sexes *in utero* electroporated with the plasmids specified in the figures and in the legends for each experiment were anaesthetized with isoflurane before decapitation. The brain was quickly removed and submerged in ice cold cutting solution containing (in mM): 110 Choline Chloride, 2.5 KCl, 1.2 NaH_2_PO_4_, 26 NaHCO_3_, 25 D-glucose, 0.5 CaCl_2_, 10 MgSO_4_. Acute 300 μm coronal slices were made of the somatosensory cortex using a vibratome (HM650V, Microm) while being saturated with 95% O_2_/5% CO_2_. The slices were immediately transferred to a submerged slice holding chamber and incubated at ±34°C for 5 min before being transferred to a second slice holding chamber also kept at ±34°C. The second holding chamber contained the same artificial cerebrospinal fluid (ACSF) as was used during all recordings and contained (in mM): 125 NaCl, 3 KCl, 1.25 NaH_2_PO_4_, 26 NaHCO_3_, 10 glucose, 2 CaCl_2_, 1 MgSO_4_. During the slicing procedure and experimental recordings, slices were saturated with 95% O_2_/5% CO_2_. Slices recovered for an hour at room temperature before starting the experiment. After the experiment, slices were fixed in 4% PFA overnight and then transferred to PBS until further processing. Whole-cell patch clamp recordings were obtained from the soma of visually identified L2/L3 pyramidal neurons from the S1 cortex with an upright microscope using IR-DIC optics (BX51WI, Olympus, Tokyo, Japan). Targeted cells in the ipsilateral side were identified by the presence of either tdTomato or GFP, depending on the experiment, elicited by an Olympus U-RFL-T burner. All recordings were done under physiological temperatures of 30± 1 °C. Patch clamp pipettes were pulled from standard wall filament borosilicate glass to obtain electrodes with a tip resistance between 2-4 MΩ. All recordings were performed using a Multiclamp 700B (Molecular Devices, Sunnyvale, CA, USA) and digitized by a Digidata 1440A (Molecular Devices, Sunnyvale, CA, USA). For the current clamp recordings, pipets were filled with a K-gluconate internal solution containing (in mM): 125 K-gluconate, 10 NaCl, 10 HEPES, 0.2 EGTA, 4.5 MgATP, 0.3 NaGTP and 10 Na-phosphocreatine. For analysis of cell morphology, biocytin (5%) was added to the intracellular solution. The final solution was adjusted to a pH of 7.2–7.4 using KOH and had an osmolarity of 280 ± 3. After getting a seal of at least 1 GΩ, whole cell configuration was obtained by applying brief negative pressure together with a short electric pulse. Prior to breaking in, cell capacitance was compensated. Series resistance was monitored but not corrected. Recordings with an unstable series resistance and higher than 20 MΩ were rejected. Membrane potentials were not corrected for liquid junction potential. Resting membrane potential was measured immediately after break in.

Each sweep started with a small and short hyperpolarizing step (−50 pA, 50 ms) to monitor access resistance. Action potentials were triggered by square step current injections into the patched neurons while holding them at −70 mV. Steps were 750 ms long and started at −300 pA with increments of 20 pA. The number of action potentials and action potential properties were analyzed using Clampfit 10.7.0.3 (Molecular Devices, LLC, USA). For each cell, the first action potential at rheobase was analyzed. The threshold was calculated by plotting the first derivative of the trace. The threshold was defined when the first derivative was lower than 10 mV/ms. Series resistance was calculated offline for each cell by plotting the difference in voltages between baseline and the hyperpolarizing steps. A linear line was plotted to visualize passive current only. The tau was calculated by fitting a standard exponential on the end of the hyperpolarizing steps. From tau and series resistance, capacitance was calculated.

### *Ex vivo* slice electrophysiology for optogenetics

P21-P25 mice of both sexes *in utero* electroporated either with the RHEBp.P37L and pCAGGS-ChR2-Venus plasmids or the empty vector and pCAGGS-ChR2-Venus plasmids (Petreanu et al., 2007), were anaesthetized with isoflurane before decapitation. The brain was quickly removed and submerged in ice cold cutting solution containing (in mM): 93 NMDG, 93 HCl, 2.5 KCl, 1.2 NaHPO_4_, 30 NaHCO_3_, 25 glucose, 20 HEPES, 5 Na-ascorbate, 3 Na-pyruvate, 2 Thiourea, 10 MgSO4, 0.5 CaCl2, 5 N-acetyl-L-Cysteine (osmolarity 310 ± 5; bubbled with 95% O_2_ / 5% CO_2_) (Ting et al., 2014). Next, 250 μm thick coronal slices were cut using a Leica vibratome (VT1000S). For the recovery, brain slices were incubated for 5 min in slicing medium at 34 ± 1°C and subsequently for ∼40 min in ACSF (containing in mM: 124 NaCl, 2.5 KCl, 1.25 Na_2_HPO_4_, 2 MgSO_4_, 2 CaCl_2_, 26 NaHCO_3_, and 20 D–glucose, osmolarity 310 ± 5mOsm; bubbled with 95% O_2_ / 5% CO_2_) at 34 ± 1°C. After recovery brain slices were stored at room temperature. For all recordings, slices were bathed in 34 ± 1°C ACSF (bubbled with 95% O_2_ / 5% CO_2_). Whole-cell patch-clamp recordings were recorded with an EPC-10 amplifier (HEKA Electronics, Lambrecht, Germany) and sampled at 20 kHz. Resting membrane potential and input resistance were recorded after whole-cell configuration was reached. Recordings were excluded if the series resistance or input resistance (RS) varied by >25% over the course of the experiment. Voltage and current clamp recordings were performed using borosilicate glass pipettes with a resistance of 3-5 MΩ that was filled with K-gluconate-based internal solution (in mM: 124 K-gluconate, 9 KCl, 10 KOH, 4 NaCl, 10 HEPES, 28.5 sucrose, 4 Na_2_ATP, 0.4 Na_3_GTP (pH 7.25-7.35; osmolarity 290 ± 5mOsm)). Recording pipettes were supplemented with 1 mg/ml biocytin to check the location of the patched cells with histological staining. Current clamp recordings were corrected offline for the calculated liquid junction potential of −10.2 mV.

Full-field optogenetic stimulation (470 nm peak excitation) was generated by the use of a TTL-pulse controlled pE2 light emitting diode (CoolLED, Andover, UK). Light intensities at 470 nm were recorded using a photometer (Newport 1830-C equipped with an 818-ST probe, Irvine, CA) at the level of the slice. To trigger neurotransmitter release from targeted axons we delivered a 1 ms light pulse with an intensity of 99.8 mW/mm^2^ at a frequency of 0.1 Hz. To ensure that we recorded action potential-driven neurotransmitter release most experiments were concluded by bath application of 10 μM tetrodotoxin (TTX), which blocked all post-synaptic responses in the recorded pyramidal neurons.

### Imaging and analysis

Images of Nissl stained sections were acquired in brightfield with a Nanozoomer scanner (Hamamatsu, Bridgewater, NJ) at a 40X resolution using the NDP.view2 software. All immunofluorescent images were acquired using a LSM700 confocal microscope (Zeiss). For the analysis of the axons *in vitro*, at least ten distinct confocal images from two different neuronal batches were taken from each coverslip for each experiment (20X objective, 0.5 zoom, 1024×1024 pixels; neurons were identified by the red immunostaining signal). The simple neurite tracer plugin from the FIJI ImageJ software was used for the analysis of the axonal length and branches. Overview images of the coronal sections were acquired by tile scan with a 10X objective. Zoom in images of the targeted area (ipsilateral) and contralateral S1 were taken using a 10X objective. For the migration analysis, confocal images (10X objective, 0.5 zoom, 1024×1024 pixels) were taken from 2 – 3 non-consecutive sections from at least 2/3 electroporated animals per condition. Images were rotated to correctly position the cortical layers, and the number of cells in different layers were counted using the ‘analyze particles’ plugin of FIJI. The results were exported to a spreadsheet for further analysis. Cortical areas from the pia to the ventricle were divided into 10 bins of equal size and the percentage of tdTomato-positive cells per bin was calculated. The soma size analysis was performed on z-stacks images acquired using a 20X objective, 1 zoom, 1024×1024 pixels, of the targeted cells in both empty vector control and *RHEB*p.P37L coronal sections. A ROI around each targeted cell in maximum intensity projection pictures was defined using the FIJI software and the area of the soma was measured using the ‘Measure’ option in ImageJ. For the analysis of pS6 intensity levels, confocal images (10X objective, 0.5 zoom, 1024×1024 pixels) of the ipsilateral and contralateral S1 cortex were acquired with the same master gain from both control and RHEB groups previously stained together against pS6 (240/244). The overall intensity level of the staining for each picture was measured using the ‘RGB measure’ plugin of FIJI and the values of each ipsilateral side were normalized against the corresponding contralateral side and plotted as averaged values. The analysis of the fluorescent intensity of the axonal branches over the contralateral cortical layers, was obtained from 3-4 matched coronal sections from at least 3 different animals per group with comparable amount of targeting. The axonal arborization was measured selecting the S1/S2 border, drawing a straight segmented line with adjusted width and length and resized in 1000 bins, and using the ‘plot profile’ option of the analyze section of FIJI to measure the fluorescent intensity of the tdTomato signal over the different layers. The values obtained for each section were exported to a spreadsheet were they were normalized against the mean background fluorescent intensity calculated on a non-targeted, cortical area of fixed size and plotted as averaged values over 10 bins of equal size. For the analysis of the morphology of biocytin filled pyramidal cells and ectopic cells in the nodule labelled with streptavidin-488 or streptavidin-Cy5, z-stacks images were taken using a 20X objective, 0.5 zoom, 1024×1024 pixels, to include the dendritic tree. Maximum intensity projection pictures were analyzed using the SynD software for the MATLAB platform to automatically detect the dendritic morphology and perform Sholl analysis (Schmitz et al., 2011).

### Statistics

Normality of the distribution for the different experiments was determined using either the Wilk– Shapiro test (Figures 1A, 1C, 2D, 3E-F, 4B, 5A, 6B, 8F and **Supplementary Figure 5B**,) or the Kolmogorov-Smirnov test (Figures 2A, 2B, 5C, 8B-C-E). Statistical analysis was performed using a one-way ANOVA (or corresponding non-parametric Kruskal-Wallis test), two-way repeated-measures ANOVA or mixed-effects analysis, Student’s t test (or corresponding non-parametric Mann-Whitney test) and correlation/association analysis. The specific test used for each experiment and relative significance are specified in the figures’ legends, in the supplementary tables or in the results section (when data are not shown in a figure). For all statistical analyses α was set at 0.05. Values are represented as average ± SEM or as median, minimum and maximum values (specified in the figures’ legends). No samples or mice were excluded from the final analysis. Group sizes, biological replicates, number of cells, samples or brain sections are indicated in the figures and their corresponding legends. All statistical tests were performed either using GraphPad Prism 8.0 (RRID: SCR_002798) or SPSS Statistics v25.0 (RRID:SCR_002865).

## ACKNOWLEDGMENTS

The authors would like to thank Charlotte de Konink, Hang Le Ha, Anne Polman and Chi Yeung for their technical contribution and Minetta Elgersma-Hooisma for managing the mouse colony. We also would like to thank Enzo Nio for advice regarding the code used to analyze the data and Steven Kushner and Dick Jaarsma for fruitful discussions. This work was supported by the Dutch TSC foundation (STSN) (G.M.v.W) and the Dutch Epilepsy foundation (Epilepsie fonds) (Y.E.).

## AUTHOR CONTRIBUTIONS

Conceptualization, M.P.O., Y.E. and G.M.v.W.; Methodology, M.P.O., Y.E. and G.M.v.W.; Investigation, M.P.O., L.M.C.K., C.B.S., M.N.; Formal Analysis: M.P.O., L.M.C.K., C.B.S., M.N.; Software, M.d.B.v.V.; Writing – Original Draft: M.P.O. and G.M.v.W.; Writing – Review & Editing: M.P.O., L.M.C.K., C.B.S., M.N., M.d.B.v.V., Z.G., Y.E., G.M.v.W.; Visualization: M.P.O.; Supervision: G.M.v.W.; Funding Acquisition: Y.E. and G.M.v.W.

