## Supplementary material for "RHEB/mTOR-hyperactivity causing cortical malformations drives seizures through increased axonal connectivity": Figures and Tables

### SUPPLEMENTAL FIGURES

#### Figure 1-figure supplement 1

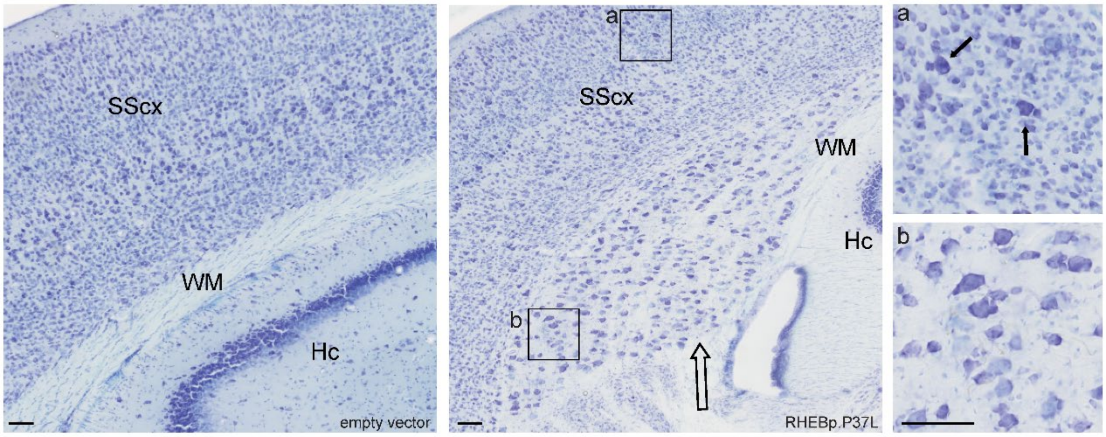


**Figure 1-figure supplement 1. Presence of heterotopia in mice *in utero* electroporated with the RHEBp.P37L construct.**

Nissl staining of coronal brain sections from 5 weeks old mice shows the presence of a clear heterotopia (indicated by the empty arrow) in the white matter (WM) of RHEBp.P37L targeted somatosensory cortex (SScx) compared to the empty vector control situation. Boxes *a* and *b* represent magnifications of layer 2 (*a*) and the heterotopia (*b*) highlighting the targeted dysplastic and enlarged cells (indicated by the arrows); Hc: hippocampus; scale bars:100 µm.

**Figure 2-figure supplement 1**

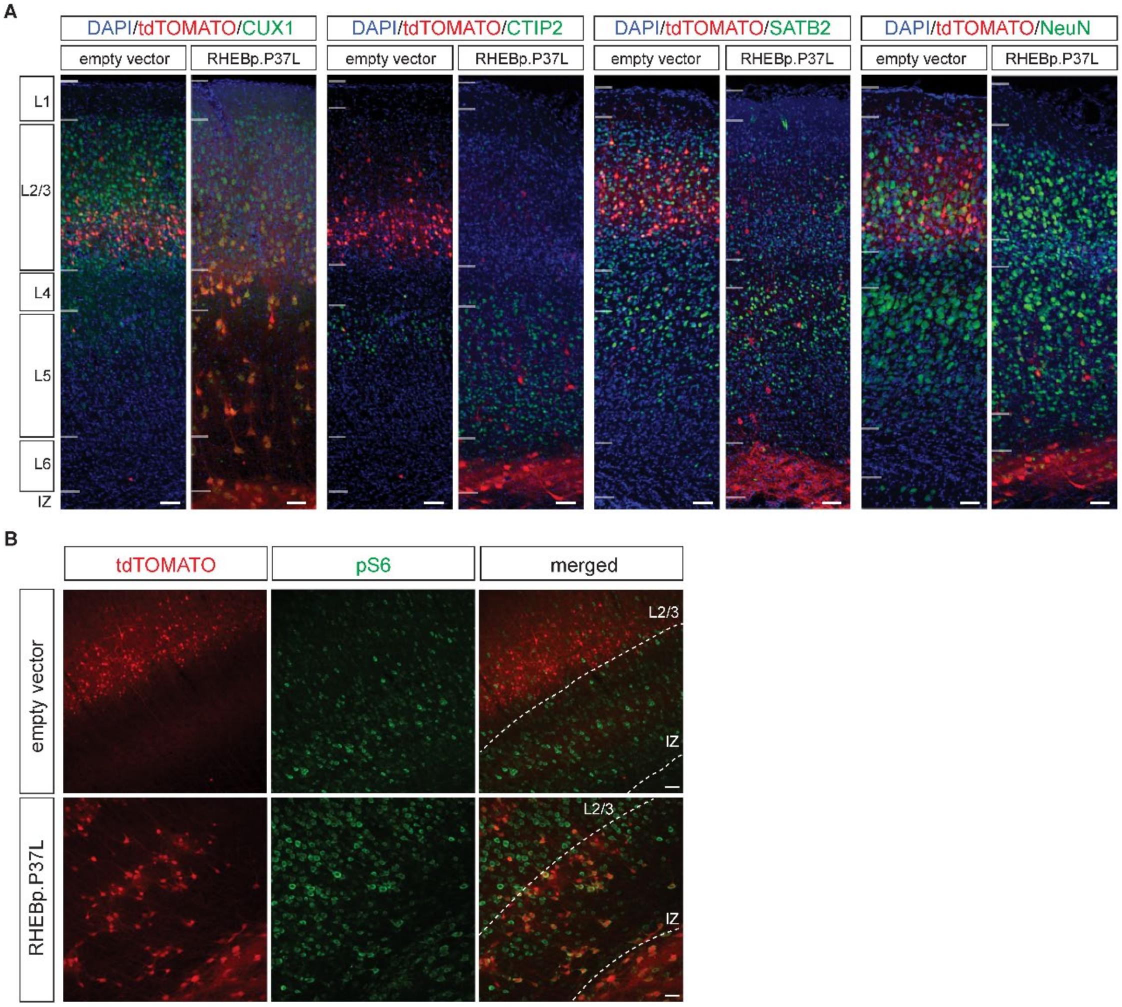


**Figure 2-figure supplement 1. Cells overexpressing the RHEBp.P37L construct maintain the molecular identity of pyramidal cells L2/3 and show mTOR hyperactivity.**

**(A)** Representative overview images of coronal sections (SScx) of empty vector control and RHEBp.P37L targeted mice (5 weeks old) probed with common cortical layers markers CUX1 (L2/3 marker), CTIP2 (L5 marker), SATB2 (cortical projection neuron marker) or NeuN (mature neuron marker). **(B)** overview of the targeted SScx of empty vector control and RHEBp.P37L targeted mice stained for pS6-240, a readout of mTOR activity; IZ: intermediate zone. Scale bars: 50 μm.

### Figure 3-figure supplement 1

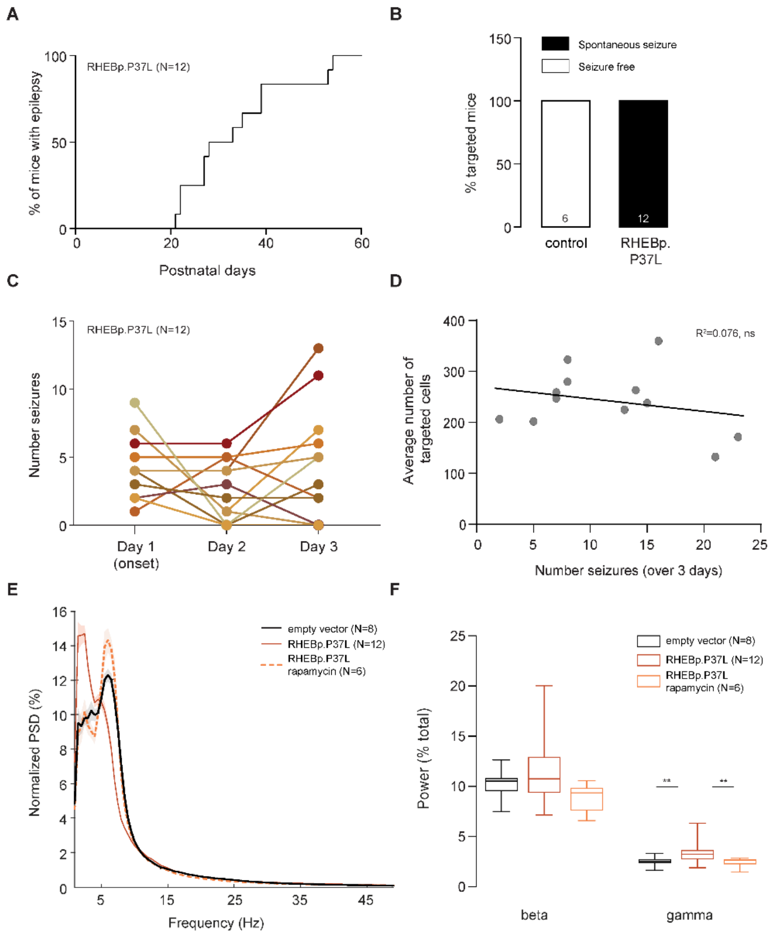


**Figure 3-figure supplement 1. RHEBp.P37L mice show spontaneous seizures that do not correlate with the number of targeted cells and alterations in the gamma frequency band**
**(A)** Onset of seizure activity for the RHEBp.P37L group (mean ± SEM: 33.33 ± 3.26; N indicates number of mice). **(B)** Percentage of targeted mice showing seizure activity measured with EEG, numbers in the bar columns indicate the number of mice per group; controls are non-targeted mice from the same litter. **(C)** Number of seizures per day for each mouse of the group in (A) and (B) across three consecutive days of recordings, with day 1 representing the onset of the very first observed seizure per mouse. **(D)** Simple scatter correlation graph with best fit regression line (Y = -2,658*X + 272,9), showing no correlation between the average number of targeted cells (measured over 3 anatomically matched non-consecutive targeted slices per mouse) and the total number of seizures over three days per animal; r(10) = -0.27, $p$ = 0.39, two-tailed Pearson's correlation; ns, non-significant. **(E)** Extended Normalized Power spectrum density (PSD) shown

in Figure 3E to include the *beta* and *gamma* frequencies (till 50 Hz); data are presented as mean (thick lines) ± SEM (shadows); N in the legend indicates number of mice per group. **(F)** Quantification of the *beta* (13-30 Hz) and *gamma* (30-50 Hz) frequency bands over the total power; box plots represent minimum and maximum value with median; N in the legend indicates number of mice per group. See **Supplemental Table 2** for statistics; ** $p<0.01$.

**Figure 5-figure supplement 1**

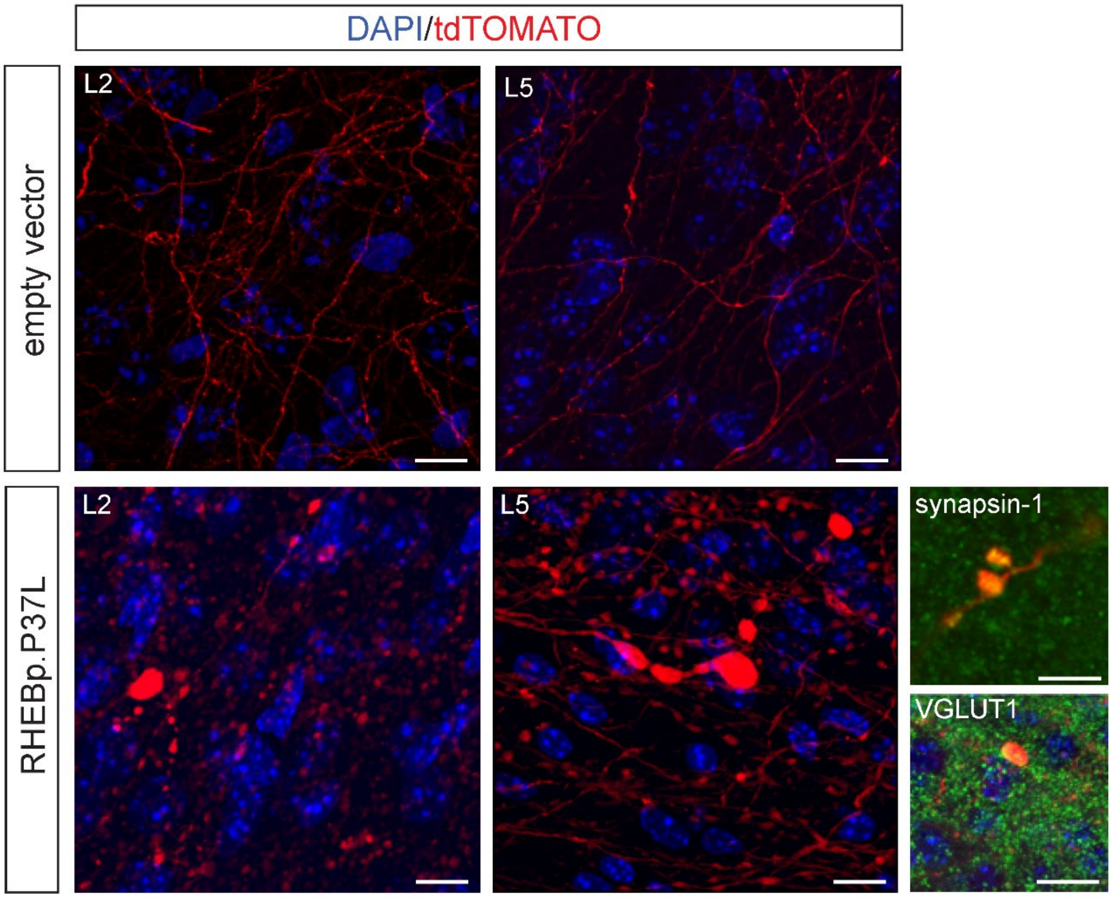


**Figure 5-figure supplement 1. Synaptic terminals in RHEBp.P37L mice have an altered morphology and are positive for Synapsin-1 and VGLUT1.**

Representative zoomed in pictures of the contralateral S1 (L2/3 and L5) of both control empty vector mice and RHEBp.P37L mice (P50); note the presence of enlarged terminals and *boutons* in RHEBp.P37L expressing cells that are positive for Synapsin-1 (a marker for synaptic vesicles, in green) and VGLUT1 (a marker for glutamatergic neurons, in green). Scale bars: 10 µm (overview), 5 µm (terminals).

**Figure 6-figure supplement 1**

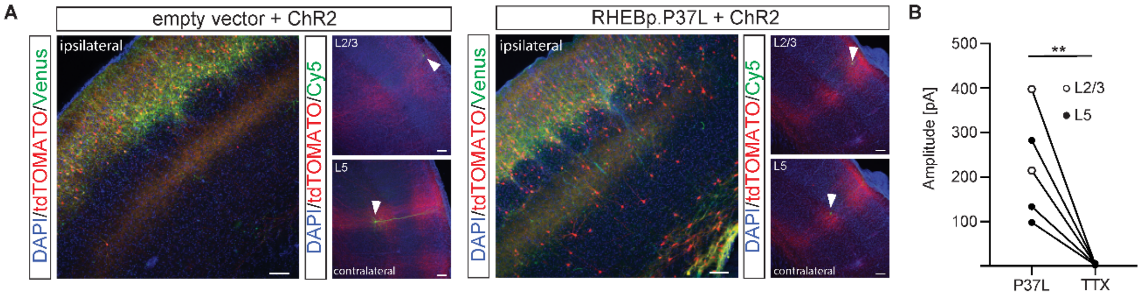


**Figure 6-figure supplement 1. Action potentials driven neurotransmitter release in RHEBp.P37L/Channelrhodopsin-2 expressing fibers.**

**(A)** Representative images showing expression of channelrhodopsin-2 (ChR2, in green) and either empty vector (left) or RHEBp.P37L (right) constructs in red (tdTomato+ cell) on the ipsilateral targeted S1; examples of contralateral patched cells in either L2/3 or L5 filled with byocitin and stained with streptavidin-Cy5 are shown for each condition and indicated with arrowheads (note that for the contralateral pictures ChR2-Venus is not shown and green represents byocitin-Cy5); scale bars: 100 µm. **(B)** Wash-in of tetrodotoxin (TTX) in RHEBp.P37L slices proves the action potential dependence of photostimulation evoked responses in L2/3 and L5; $t(5)=4.8$, $p=0.005$, two-tailed paired t-test; ** $p<0.01$.

**Figure 8-figure supplement 1**

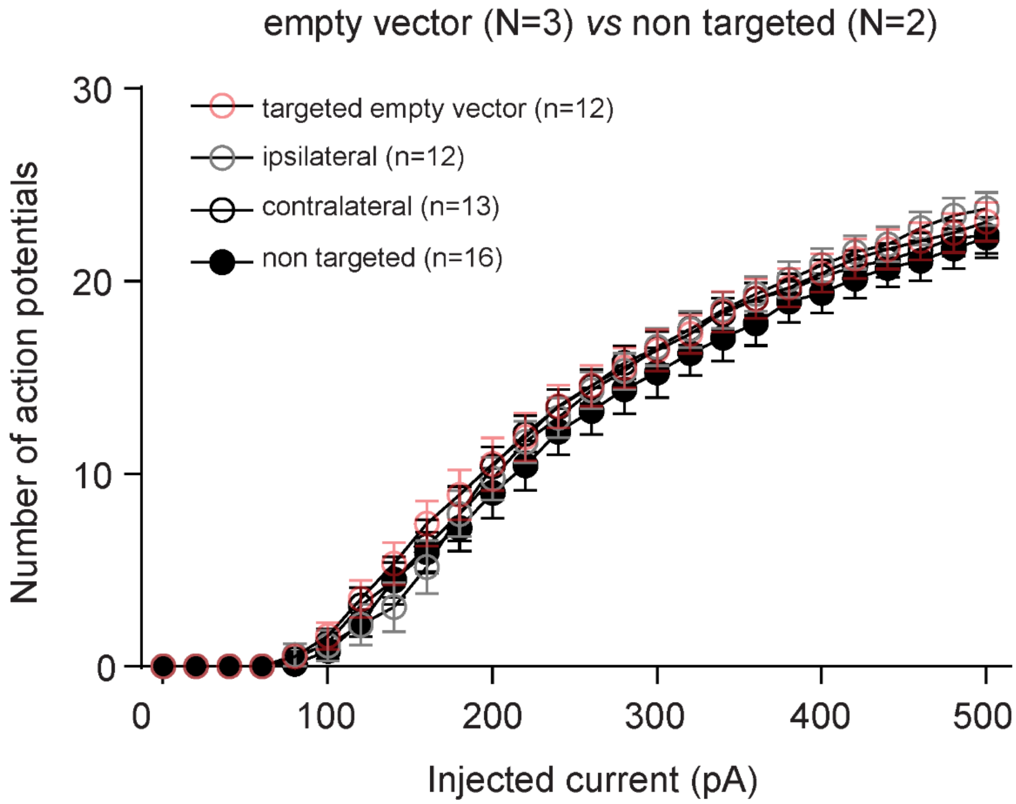


**Figure 8-figure supplement 1. Excitability phenotype of control cells from empty vector control and non-targeted mice.**

Number of action potentials in response to increasing depolarizing currents shows that there's no difference in excitability in empty vector targeted mice or non-targeted mice; data are presented as mean ± SEM; interaction injected current/group condition: F (75, 1225) = 0.7275, non-significant; mixed-effects analysis; N = number of mice and n = number of cells analyzed.

**SUPPLEMENTARY TABLES**

| **Supplementary Table 1. Statistical analysis related to Figure 1A** | | | |
|---|---|---|---|
| **Test applied: Two-way ANOVA** | | | |
| **Source of variation** | **F (DFn, DFd)** | **P value** | **P value summary** |
| TSC -/+ | F (1, 35) = 13.38 | 0.0008 | *** |
| Group condition | F (3, 35) = 42.39 | <0.0001 | **** |
| Interaction | F (3, 35) = 2.377 | 0.0866 | ns |
| **Post hoc: Sidak's multiple comparisons test** | | | |
| | **Comparison** | **Adjusted P value** | **P value summary** |
| TSC – *vs* TSC+ | RHEB wt | 0.0486 | * |
| | RHEBp.S16H | 0.0049 | ** |
| | RHEBp.P37L | 0.9791 | ns |
| | Empty vector control | 0.9621 | ns |
| **Post hoc: Tukey's multiple comparisons test** | | | |
| | **Comparison** | **Adjusted P value** | **P value summary** |
| TSC – | RHEB wt vs RHEBp.S16H | 0.0205 | * |
| | RHEB wt vs RHEBp.P37L | 0.0001 | *** |
| | RHEB wt vs Empty vector control | 0.0151 | * |
| | RHEBp.S16H vs RHEBp.P37L | 0.2859 | ns |
| | RHEBp.S16H vs Empty vector control | <0.0001 | **** |
| | RHEBp.P37L vs Empty vector control | <0.0001 | **** |
| TSC+ | RHEB wt vs RHEBp.S16H | 0.3696 | ns |
| | RHEB wt vs RHEBp.P37L | <0.0001 | **** |
| | RHEB wt vs Empty vector control | 0.7682 | ns |
| | RHEBp.S16H vs RHEBp.P37L | 0.0009 | *** |
| | RHEBp.S16H vs Empty vector control | 0.0603 | ns |
| | RHEBp.P37L vs Empty vector control | <0.0001 | **** |

ns: non-significant, * $p<0.05$, ** $p<0.01$, *** $p<0.001$, **** $p<0.0001$

| | | | |
|---|---|---|---|
| **Supplementary Table 2. Statistical analysis related to Figures 3E-F and Figure 3 – figure supplement 1** | | | |
| **Test applied LFP: Two-way RM ANOVA** | | | |
| **Source of variation** | **F (DFn, DFd)** | **P value** | **P value summary** |
| Band Frequency | F (1.100, 53.92) = 353.7 | P<0.0001 | **** |
| Group condition | F (2, 49) = 1.011 | P=0.3714 | ns |
| Interaction Band frequencies/ group condition | F (6, 147) = 7.456 | <0.0001 | **** |
| **Tukey's multiple comparisons test** | **Comparison** | **Adjusted P value** | **P value summary** |
| delta | empty vector control vs. RHEBp.P37L | 0.0479 | * |
| | empty vector control vs. RHEBp.P37L - rapamycin | 0.9614 | ns |
| | RHEBp.P37L vs. RHEBp.P37L - rapamycin | 0.0856 | ns |
| theta | empty vector control vs. RHEBp.P37L | 0.0024 | ** |
| | empty vector control vs. RHEBp.P37L - rapamycin | 0.8288 | ns |
| | RHEBp.P37L vs. RHEBp.P37L - rapamycin | 0.0053 | ** |
| beta | empty vector control vs. RHEBp.P37L | 0.2043 | ns |
| | empty vector control vs. RHEBp.P37L - rapamycin | 0.0637 | ns |
| | RHEBp.P37L vs. RHEBp.P37L - rapamycin | 0.0071 | ** |
| gamma | empty vector control vs. RHEBp.P37L | 0.0039 | ** |
| | empty vector control vs. RHEBp.P37L | 0.8465 | ns |
| | empty vector control vs. RHEBp.P37L - rapamycin | 0.0016 | ** |
| **Test applied ratio theta/delta: Kruskal-Wallis test** | | | |
| **Kruskal-Wallis H** | **df** | **P value** | **P value summary** |
| 13.37 | 2 | 0.0012 | ** |
| **Dunn's multiple comparisons test** | **Comparison** | **Adjusted P value** | **P value summary** |
| ratio theta/delta | empty vector control vs. RHEBp.P37L | 0.0169 | * |
| | empty vector control vs. RHEBp.P37L - rapamycin | >0.9999 | ns |
| | RHEBp.P37L vs. RHEBp.P37L - rapamycin | 0.0036 | ** |

ns: non-significant, * $p<0.05$, ** $p<0.01$, **** $p<0.0001$

| | | | | |
|---|---|---|---|---|
| **Supplementary Table 3. Statistical analysis related to Figure 6** | | | | |
| **Test applied: Two-tailed Mann-Whitney test** | | | | |
| **Parameter/ Layer** | **Unit** | **Mean ± SEM** | **Mann-Whitney U P value** | **P value summary** |
| Amplitude L2/3 | pA | control: 54.54±10.77 RHEBp.P37L: 356.80±54.06 | U = 7 p=0.0004 | *** |
| Amplitude L5 | pA | control: 132.0±45.71 RHEBp.P37L: 365.70±97.59 | U = 34 p=0.0226 | * |
| Charge L2/3 | pA*ms | control: 353.40±82.25 RHEBp.P37L: 3079.0±516.90 | U = 6 p=0.0003 | *** |
| Charge L5 | pA*ms | control: 1552.0±816.10 RHEBp.P37L: 3658.0±1007.0 | U = 36 p=0.0308 | * |
| Vm L2/3 | mV | control: -77.83±1.50 RHEBp.P37L: -76.43±2.22 | U = 51.50 p=0.9851 | ns |
| Vm L5 | mV | control: -73.0±0.55 RHEBp.P37L: -73.53±1.16 | U = 70 p=0.7933 | ns |
| R series L2/3 | MΩ | control: 11.97±0.94 RHEBp.P37L: 11.88±0.71 | U = 51 p=0.9576 | ns |
| R series L5 | MΩ | control: 14.91±1.11 RHEBp.P37L: 12.46±0.86 | U = 47 p=0.1289 | ns |
| Rm L2/3 | MΩ | control: 105.50±10.92 RHEBp.P37L: 97.08±8.69 | U = 45 p=0.6324 | ns |
| Rm L5 | MΩ | control: 131.50±22.75 RHEBp.P37L: 130.70±8.22 | U = 57 p=0.3383 | ns |

ns: non-significant, * $p$<0.05, *** $p$<0.001

**Supplementary Table 4. Seizure frequency in adult RHEBp.P37L/LSL-TeTxLC mice upon tamoxifen administration related to Figures 7C-7D**

| **Number seizures/24hr** | **Mouse 1** | **Mouse 2** | **Mouse 3** | **Mouse 4** | **Mouse 5** | **Mouse 6** |
|---|---|---|---|---|---|---|
| Day 0 (seizures onset) | 5 | 5 | 7 | 6 | 6 | 5 |
| Day 1 - Tamoxifen | 1 | 8 | 6 | 7 | 6 | 1 |
| Day 2 - Tamoxifen | 0 | 6 | 3 | 0 | 3 | 2 |
| Day 3 - Tamoxifen | 0 | 0 | 1 | 0 | 0 | 1 |
| Day 4 - Tamoxifen | 0 | 0 | 2 | 0 | 0 | 0 |
| Day 5 | 0 | 0 | 0 | 0 | 0 | 2 |
| Day 6 | 0 | 0 | 0 | 0 | 0 | 0 |
| Day 7 | 0 | 0 | 0 | 0 | 3 | 1 |
| Day 8 | - | 0 | 0 | 0 | 0 | 0 |
| Day 9 | - | 0 | 0 | 0 | 2 | 2 |
| Day 10 | - | - | - | - | 0 | - |
| Day 24 | 0 | 0 | 0 | 0 | 0 | 0 |
| Day 25 | 0 | 0 | 0 | 0 | 0 | 0 |
| Day 26 | 0 | 0 | 0 | 0 | 0 | 0 |
| Day 44 | 0 | 0 | 0 | 0 | 0 | 0 |
| Day 45 | 0 | 0 | 0 | 0 | 0 | 0 |

**Supplementary Table 5. Statistical analysis related to Figure 8B-C**

**Test applied on basic properties: One-way ANOVA**

| Basic properties | F (DFn, DFd) | P value | P value summary |
|---|---|---|---|
| Cm | F (3, 111) = 6.525 | P=0.0004 | *** |
| Rm | F (3, 111) = 10.47 | P<0.0001 | **** |
| Vm | F (3, 111) = 0.3580 | P=0.7834 | ns |

**Post hoc: Tukey's multiple comparisons test**

| Comparison | Cm Adjusted P Value | Cm Summary | Rm Adjusted P Value | Rm Summary | Vm Adjusted P Value | Vm Summary |
|---|---|---|---|---|---|---|
| control vs. contralateral RHEBp.P37L | 0.9218 | ns | 0.9970 | ns | 0.9028 | ns |
| control vs. ipsilateral RHEBp.P37L | 0.1568 | ns | 0.6241 | ns | 0.9974 | ns |
| control vs. targeted RHEBp.P37L | 0.0003 | *** | <0.0001 | **** | 0.9585 | ns |
| contralateral RHEBp.P37L vs. ipsilateral RHEBp.P37L | 0.6314 | ns | 0.6506 | ns | 0.9751 | ns |
| contralateral RHEBp.P37L vs. targeted RHEBp.P37L | 0.0171 | * | <0.0001 | **** | 0.7394 | ns |
| ipsilateral RHEBp.P37L vs. targeted RHEBp.P37L | 0.2425 | ns | 0.0033 | ** | 0.9337 | ns |

**Test applied on excitability (RHEBp.P37L vs control): Mixed-effects model analysis**

| Source of variation | F (DFn, DFd) | P value | P value summary |
|---|---|---|---|
| Injected current | F (2,770, 265.6) = 1228 | <0.0001 | **** |
| Group condition | F (3, 96) = 44.64 | <0.0001 | **** |
| Interaction current/condition | F (75, 2397) = 29.02 | <0.0001 | **** |

**Post hoc: Tukey's multiple comparisons test**

| Main effect: group condition | Mean difference | Adjusted P Value | P value summary |
|---|---|---|---|
| targeted RHEBp.P37L vs. ipsilateral RHEBp.P37L | 7.349 | <0.0001 | **** |
| targeted RHEBp.P37L vs. contralateral RHEBp.P37L | -0.2674 | <0.0001 | **** |
| targeted RHEBp.P37L vs. control | -2.948 | <0.0001 | **** |
| ipsilateral RHEBp.P37L vs. contralateral RHEBp.P37L | -7.616 | <0.0001 | **** |
| ipsilateral RHEBp.P37L vs. control | -10.3 | 0.9447 | ns |
| contralateral RHEBp.P37L vs. control | -2.681 | <0.0001 | **** |

ns: non-significant, * $p$<0.05, ** $p$<0.01, *** $p$<0.001, **** $p$<0.0001

**Supplementary Table 6. Statistical analysis related to Figure 8E-F**

**Test applied on basic properties: One-way ANOVA**

| **Basic properties** | **F (DFn, DFd)** | **P value** | **P value summary** |
|---|---:|---:|---:|
| Cm | F (3, 69) = 10.43 | P<0.0001 | **** |
| Rm | F (3, 69) = 16.01 | P<0.0001 | **** |
| Vm | F (3, 73) = 0.8526 | P=0.4697 | ns |

**Post hoc: Tukey's multiple comparisons test**

| **Comparison** | **Cm Adjusted P Value** | **Cm Summary** | **Rm Adjusted P Value** | **Rm Summary** | **Vm Adjusted P Value** | **Vm Summary** |
|---|---:|---:|---:|---:|---:|---:|
| targeted RHEBp.P37L vs. targeted RHEBp.P37L/LSL-TeTxLC | 0.1347 | ns | 0.7179 | ns | 0.6715 | ns |
| targeted RHEBp.P37L vs. contralateral RHEBp.P37L/LSL-TeTxLC | 0.0087 | ** | 0.0004 | *** | 0.9996 | ns |
| targeted RHEBp.P37L/LSL-TeTxLC vs. contralateral RHEBp.P37L/LSL-TeTxLC | <0.0001 | **** | <0.0001 | **** | 0.7032 | ns |
| targeted RHEBp.P37L/LSL-TeTxLC vs. contralateral RHEBp.P37L | 0.0006 | *** | <0.0001 | **** | 0.6425 | ns |
| contralateral RHEBp.P37L/LSL-TeTxLC vs. contralateral RHEBp.P37L | 0.5782 | ns | 0.9675 | ns | 0.9971 | ns |
| targeted RHEBp.P37L vs. targeted RHEBp.P37L/LSL-TeTxLC | 0.1347 | ns | 0.7179 | ns | 0.6734 | ns |

**Test applied on excitability: Mixed-effects model analysis**

| **Excitability RHEBp.P37L/ LSL-TeTxLC** | **F (DFn, DFd)** | **P value** | **P value summary** |
|---|---|---|---|
| Injected current | F (2.357, 251.5) = 870.7 | <0.0001 | **** |
| Group condition | F (4, 107) = 3714 | <0.0001 | **** |
| Interaction current/condition | F (100, 2667) = 21.99 | <0.0001 | **** |

**Post hoc: Tukey's multiple comparisons test**

| **Comparison** | **Mean difference** | **Adjusted P Value** | **P value summary** |
|---|---|---|---|
| control vs. targeted RHEBp.P37L/LSL-TeTxLC | 6.556 | <0.0001 | **** |
| control vs. contralateral RHEBp.P37L/LSL-TeTxLC | -0.3674 | 0.9490 | ns |
| targeted RHEBp.P37L vs. targeted RHEBp.P37L/LSL-TeTxLC | -0.7935 | 0.4371 | ns |
| targeted RHEBp.P37L vs. contralateral RHEBp.P37L/LSL-TeTxLC | -7.717 | <0.0001 | **** |
| contralateral RHEBp.P37L vs. targeted RHEBp.P37L/LSL-TeTxLC | 9.504 | <0.0001 | **** |
| contralateral RHEBp.P37L vs. contralateral RHEBp.P37L/LSL-TeTxLC | 2.581 | 0.0002 | *** |

ns: non-significant, ** $p<0.01$, *** $p<0.001$, **** $p<0.0001$
